# Cautious explorers: comparing movement patterns of wild and rewilded solitary predators

**DOI:** 10.1101/2025.10.15.682496

**Authors:** Mohammad S. Farhadinia, Viatcheslav V. Rozhnov, Luciano Atzeni, Jose A. Hernandez-Blanco, Anna A. Yachmennikova, Maria D. Chistopolova, Alexander N. Minaev, Natalia A. Dronova, Alim B. Pkhitikov, Kaveh Hobeali, Pavel I. Weinberg, Sergey A. Trepet, Paul J. Johnson, David W. Macdonald

## Abstract

Many rewilding projects emphasize recovering trophic networks through species translocation. Understanding the behaviour and movement patterns of rewilded animals is central to evaluating the success of rewilding efforts. In this study, we compared the behaviour of rewilded Persian leopards (*Panthera pardus tulliana*) in Russia with their wild counterparts in Iran to assess how they compare in terms of exploratory and cautious movement in new environments. We used Behavioural Change Point Analysis with selection procedures to identify points where individuals switched between movement modes. We analysed intrinsic movement metrics (speed, persistence velocity, turning velocity) and spatiotemporal attributes (duration, net squared displacement, distance travelled). Time spent in each behavioural mode differed between wild and rewilded leopards: on average, rewilded leopards spent more time in the “Ranging” mode and less time “Encamped” compared to wild leopards. Rewilded leopards were also slower than wild leopards, particularly in the “Encamped” state while wild leopards exhibited similar speeds across modes. Importantly, rewilded leopards had larger displacement and spent longer in the “Ranging” mode, a pattern not seen in wild leopards. These findings highlight distinct spatial behaviour patterns between wild and rewilded leopards. Accordingly, rewilded leopards adopted both exploratory and cautious behavioural strategies, reflecting their tendency to range more widely over extended periods, plausibly to gather information from large parts of their environments, albeit at lower speed, arguably to avoid potential risks, suggesting that they are in the process of adapting to new environments. These differences are likely to reflect different ecological circumstances, but even more so the previous experiences and thus different social circumstances of the two samples of Persian leopards.

**OPEN RESEARCH STATEMENT:** All locational data from Iran are publicly available on Movebank: https://www.movebank.org/panel_embedded_movebank_webapp. Project: Persian leopard Tandoureh Iran (accession number 270329098). All the data are available for download as .csv file.

## INTRODUCTION

Conservation translocations, where animals are deliberately moved to achieve a conservation net gain, are widely used to re-establish extinct populations or to reinforce an existing population of conspecifics (IUCN 2013). Movement is the first behavioural response of translocated animals to “forced dispersal” in a new environment (Stamps & Swaisgood, 2007). Importantly, the development of stable home ranges depends on spatial learning where individuals acquire information necessary for satisfying basic needs such as food acquisition, escape from predators and competitors, and access to reproduction sites (Sulikowski and Burke 2011). This information also affects decisions to change movement patterns (e.g., switching to area-restricted search in regions of high forage availability) (Börger et al., 2008; Lewis et al., 2021). Therefore, understanding the post-release movement trajectories of translocated animals is central for exploring how animals develop spatial memory (Falcón-Cortés et al. 2021, Ranc et al. 2022) and establish their home ranges (Börger et al. 2008, Van Moorter et al. 2009) which eventually contributes to the success of such initiatives.

Understanding the development of spatial memory can help practitioners to optimize rewilding initiatives. Defined as the creation of ecological interactions to promote self-regulating biodiverse ecosystems, particularly through the recovery of trophic networks (Svenning et al. 2016). Globally, there is a general emphasis on rewilding associated with the announcement of the 2020s as the Decade of Ecosystem Restoration by the United Nations (Cooke et al. 2019). Inspired by this political attention, improvements in carnivore translocation have the potential to make a substantial contribution to nature recovery (Thomas et al. 2023).

An individual’s immediate response after introduction to a novel environment is first physiological, followed by movement; together these shape its behavioural plasticity. This plasticity allows an animal to adjust its behaviour to suit the conditions of its immediate environment under altered conditions and, in so doing, increase its fitness (Wong and Candolin 2015). Certain parts of the environment retain biologically important information over time, forming what was initially described as the “biological signal field” as external trigger of animal movement (Naumov 1973).

Equally important, movement is a complicated behavioural process that depends on an animal’s internal state, its physiological constraints, and its environment, the outcome of which is a movement path consisting of a mixture of different movement modes (Gurarie et al. 2009). These various behaviours should leave detectable signatures in movement paths; decomposing that mixture into its component parts is a challenging task but nonetheless technically feasible (Gurarie et al. 2009, Falcón-Cortés et al. 2021).

Trajectory segmentation represents a set of methods, where variation in intrinsic movement metrics, such as speed, step length and turning angle is used to identify segments of behaviour (i.e. modes) characterised by shared characteristics from which it is possible to infer behavioural information (Edelhoff et al. 2016, Soleymani et al. 2017). Gurarie et al. (2009) developed a phenomenological time-series method to conduct behavioural change point analysis (BCPA) for identifying the structure or periodicity of movement paths adapting techniques with origins in time series and signal processing while accounting for autocorrelated and irregularly sampled data. This method used persistence velocity (the tendency of a movement to persist in a particular direction), turning velocity (the tendency of a movement to turn in a perpendicular direction) and Net Squared Displacement (NSD) (Gurarie et al., 2009), to provide consistent metrics of movement applicable to dispersing wide-ranging carnivores such as wolf (*Canis lupus*) (Gurarie et al. 2016) and puma (*Puma concolor*) (Gigliotti et al. 2019).

In this study, with the ultimate objective of parsing out the spatio-temporal movement parameters characteristic of translocated leopards, and thereby of informing reintroduction protocols, we compared wild Persian leopards *(Panthera pardus tulliana)* from the Iran-Turkmenistan borderlands with captive-born leopards of the same subspecies reintroduced to the Russian Caucasus. The reintroduction project, initiated in 2007, aimed to establish two Persian leopard population nuclei in the northern part of their historical range, where they had disappeared since the 1950s. Despite the first releases in 2016, no breeding has been observed in Russia as of September 2024 (Rozhnov et al. 2022). Clearly, our two samples of Persian leopards differed in several respects, of which one category was related to their provenances and previous experiences (and thus crucial to the success of rewilding), whereas the second category concerns differences in ecological circumstances between the study sites. Our parsing out of the former category depends on correctly identifying, and thus taking account of, the ecological differences. We elaborate this argument below, but first we identify the two hypotheses we will evaluate with respect to two distinct behavioural strategies:

H1) Exploratory strategy hypothesis: Given the lack of familiarity with the new environment, we hypothesised that rewilded leopards show more exploratory pattern by spending more time in highly mobile modes, perhaps motivated to acquire information, and these movement modes would be generally longer (both in time and distance) than those of wild leopards. Conversely, we expected that wild leopards would generally spend more time in less mobile modes, cover longer distance in this mode and make longer excursions, paralleling observations of other felids (Weise et al. 2015, Yiu et al. 2015, Sievert et al. 2022).

H2) Cautious strategy hypothesis: Rewilded leopards exhibit cautious behaviour, characterized by slower and less direct movements, often associated with “shy” personality traits (Spiegel et al. 2017). This cautious behaviour may improve survival during the critical early stages of reintroduction when the animal has minimal knowledge about its environment, and the unpredictability of that environment is at its highest level for the animal (Popov 2010). In contrast, wild leopards display faster and more direct movements, similar to other translocated felids (Cisneros-Araujo et al. 2024).

## MATERIALS AND METHODS

### Ethics statement

The Iranian Department of Environment reviewed all sampling, trapping and handling procedures and approved permits for the work conducted in Iran (93/16270). The trapping and handling protocol for activities in Iran was also approved by the University of Oxford’s Ethical Review Committee (BMS-ERC-160614). In Russia, the leopards were handled in accordance with the ethical protocol #_33a approved on 08.10.2019, extended on 07.07.2022 and signed by the Biological Ethical Review Commission of the A.N.Severtsov Institute of Ecology and Evolution Russian Academy of Sciences.

### Study area

The current study was conducted within three sites (Figure 1 & S1; Table 1):

**Figure 1.**
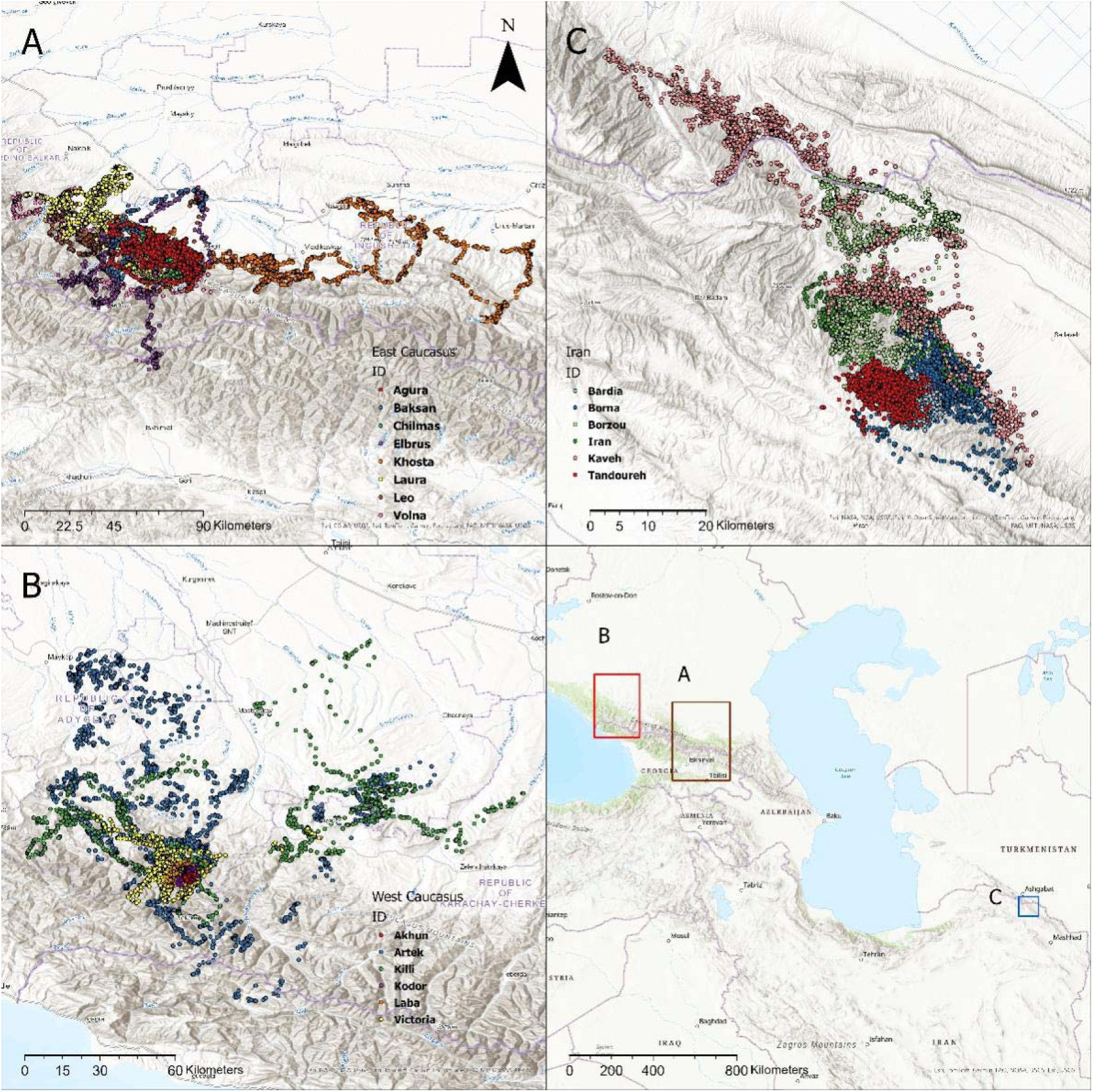
Study area overview representing collared leopard fixes in the A) Central Caucasus, Russia; B) Western Caucasus, Russia, C) Tandoureh National Park, Iran.

**Table 1.** Details of study areas in Iran and Russia.

| **Study area** | **Size**<br>**(km<sup>2</sup>)** | **Number of collared**<br>**leopards (M:F)** | **Wild prey species** | **Other large**<br>**carnivores** |
| --- | --- | --- | --- | --- |
| Tandoureh National Park, Iran | 355 | 6 (5:1) | Urial, bezoar goat and wild pig | Grey wolf and Eurasian lynx (both occasionally) |
| Caucasian Nature Biosphere Reserve, Western Caucasus, Russia | 2,800 | 6 (4:2) | Wild pig, roe deer, red deer, chamois, Caucasian tur and European bison | Grey wolf, brown bear and Eurasian lynx |
| Alaniya National Park, North-Ossetian Nature Reserve, Turmonskiy Wildlife Sanctuary and adjacent lands, Central Caucasus, Russia | 7,800 | 8 (4:4) | Wild pig, roe deer, red deer, chamois, Caucasian tur, raccoon dog, European badger, golden jackal, red fox and wildcat | Grey wolf, brown bear and Eurasian lynx |

In Iran, we studied leopards in Tandoureh National Park, north-eastern Iran, starting in September 2014. The park, protected since 1968, spans 355 km² with elevations from 1,000 to 2,600 meters and annual precipitation of 250 to 300 mm. It is characterized by mountains covered with wormwood *Artemisia* sp. and scattered juniper trees *Juniperus* sp. Local communities, mainly sheep and goat herders, live in nearby outside villages. Wild prey is available only inside the park, except wild pigs, which are occasionally found in multi-use areas, outside the national park.

In Russia, we tracked rewilded leopards in two areas: the Western Caucasus and the Central Caucasus. The Western Caucasus, extending over the extreme western end of the Caucasus mountains, is one of the few large mountain areas of Europe that has not experienced significant human impact. Its subalpine and alpine pastures have only been grazed by wild animals, with extensive tracts of undisturbed mountain forests. Leopards were released in the Federal Caucasian Nature Reserve with an area of 2,800 km^2^ which is the western part of the Greater Caucasus.

The Central Caucasus is situated around 380 km to the east from the Western Caucasus within the Republic of Kabardino-Balkaria, the Republic of North Ossetia-Alania, and the Chechen Republic, which all are situated in the northern Greater Caucasian Ridge along the border of the Russian Federation and Georgia. The geographical characteristic of the region describes the middle mountains, the transitional Alpine zone and highlands. The mid-mountain area, which ranges from 1,000 to 1,800 meters above sea level, is covered mainly by broad-leaved forests and rock massifs with an admixture of conifers on the northern slopes while shrubs and alpine zones are seen in higher altitudes ((Rozhnov and Lukarevsky 2008).

Although ecological differences exist between the three study areas, they share a wide range of features. These include predominantly montane landscapes with significant altitude variability, severe winters with snow-covered ridgelines, moderately similar intraguild compositions (except for the presence of brown bears in the Russian study sites), and extremely low human density within the reserves. Therefore, we expect that the ecological differences are sufficiently minor to only explain a small proportion of the differences in resident behaviour between the areas, while their origin should play a major role in shaping their behaviours.

#### Leopard handling and collaring

In Iran, leopards were captured with modified Aldrich foot-snares, monitored every 1-2 hours using VHF transmitters (Wildlife Materials, Inc., Illinois, USA; see Farhadinia et al. (2017b) for more details). Leopards were fitted with GPS collars (LOTEK Engineering Ltd., ON, Canada).

In Russia, leopards from Sochi Breeding Centre (SBC) were released into two study sites (Table S1) after training in captivity (see (Hernandez-Blanco et al., (2024); Rozhnov et al., (2020)). They were immobilized to fit GPS collars (heterogeneous mix) with fixes programmed for up to 24 times per day. Transported by helicopter, the leopards were released directly into the wild (i.e. no ‘soft release’ involving an adaptation enclosure). Out of 14 leopards released, 12 survived until 6 months post-release, and 8 were still alive by the end of their first year (Table S1; Figure S1; see Supplementary Information for further details).

### Statistical analysis

All analyses were performed in the R statistical environment version (R Development Core Team 2013).

#### Data screening

We omitted the first 4 days of GPS data for Iranian leopards to minimise the influence of any abnormal behaviour post-collaring. GPS fixes were inspected for errors, outliers, and unrealistic movement patterns using a predefined distance threshold and speed parameters (Bjørneraas et al. 2010). Erroneous data points were removed through the ‘atlastools’ and ‘ctmm’ package (Calabrese et al. 2016, Gupte et al. 2022) (see Supplementary Information and Table S2 for further details).

#### Path segmentation to classify movement modes

We interpolated tracks at regular two hourly intervals using the ‘waddle’ package (Gurarie et al. 2016). Our analysis involved a three-step process. First, the tracks were split into segments using the BCPA framework (Gurarie et al. 2009, 2016). The ‘bcpa’ package (Gurarie 2014) was used with persistence velocity (speed multiplied by the cosine of the turning angle, i.e., V*cosθ as the key response variable (Gurarie et al. 2009). This metric is crucial as it quantifies both the direction and intensity of an animal’s movement tendency (Gurarie 2014). Second, linear and non-linear equations were identified to describe potential candidate movement modes (Bunnefeld et al. 2011, Oswald et al. 2012). Third, the segments were then classified based on the fit of these equations to the patterns of NSD (Börger and Fryxell 2012) for each segment, calculated from the origin to the end of each segment.

In BCPA, each response variable is modelled as a Gaussian process, characterized by parameters such as mean, standard deviation, and autocorrelation. The method identifies significant changes in the autocorrelation parameter, both abrupt and gradual, by employing a technique that sweeps across the time series data with analysis windows, selecting the optimal window based on the Bayesian Information Criterion. The BCPA approach does not require any prior assumptions about the number of segments or states within the data (Gurarie et al. 2009, 2016).

User-defined parameters within BCPA include the moving window size, which determines the temporal scale for identifying meaningful biological changes; the window step size, which specifies the incremental shifts of the analysis window and affects processing speed; and the *K* sensitivity parameter, designed to adjust for the window size and prevent the detection of false change points. Given the absence of predefined expectations for these parameters, a comprehensive sensitivity analysis was undertaken. This involved varying the window size (20, 30, 40, 50), window step (1, 2, 3, 4), and K value (1, 2, 3, 4), culminating in 64 distinct BCPA models per animal and a total of 1,280 models.

To determine the most effective parameter combination for each variable, the residual of the models and the distribution’s adherence to normality was assessed with the Anscombe-Glynn kurtosis test in the ‘moments’ package (Komsta and Novomestky 2015). The aim was to identify parameter settings that yielded a kurtosis test value closest to 3, indicative of a perfect normal distribution and thus minimal excess kurtosis, for each variable and individual.

We then adopted the set of candidate models developed by Morelle et al. (2017) to describe movement modes in leopards. Specifically, we hypothesised five candidate modes: 1) ‘Encamped’ (affinity toward a certain area and undirected movement), 2) ‘Wandering’ (non-directional movement leading to a linear increase in displacement), 3) ‘Ranging’ (directional movement from an affinity area leading to an increase in the displacement followed by a stabilisation, i.e., the arrival to another affinity area or returning to the original areas), 4 and 5) ‘round-trips’ (full and partial, indicating displacement farther from the origin with return to the starting or a close-by location, respectively; Figure S2).

Our mode classification relies on applying both linear and non-linear equations to the shape of the NSD for each segment, calculated using the ‘adehabitatLT’ package (Calenge 2006). We distinguished between ‘Encamped’ and ‘Wandering’ modes using linear equations, and ‘Full Round-trip’, ‘Partial Round-trip’, and ‘Ranging’ modes with non-linear equations (Figure S2), following the precedents in previous work (Bunnefeld et al. 2011, Singh et al. 2012, Morelle et al. 2017). We used the ‘FlexParamCurve’ package (Oswald et al. 2012) to fit distinct types of non-linear equations. Linear equations were derived through ordinary least squares regression. For non-linear models, we used the ‘FlexParamCurve’ package to initialise suitable starting parameters for the curves.

To evaluate the goodness-of-fit for each candidate movement mode per segment, we employed the Concordance Criterion (CC) (Singh et al. 2012). This measure, particularly useful for non-linear models, quantifies the agreement level between empirical and predicted values on a scale from –1 (complete negative fit) to 1 (complete positive fit). However, the CC is not suitable for assessing the ‘Encamped’ mode (intercept-only models), for which the Akaike Information Criterion (AIC) is predominantly used.

The classification into movement modes primarily depends on attaining the highest CC value (≥ 0.7 for ‘Ranging’ and ‘Round-trip’ modes) or specific conditions for identifying the ‘Wandering’ mode, should the CC be ≥ 0.4. ‘Encamped’ mode is confirmed when it secures the lowest AIC, if the highest CC is < 0.7 for all modes except ‘Wandering’ (where CC is < 0.4), or when ‘Wandering’ registers the highest CC yet remains < 0.4. ‘Wandering’ is recognised when it attains the highest CC (assuming CC ≥ 0.4) or if ‘Ranging’ and ‘Round-trip’ modes do not meet the 0.7 CC threshold, provided ‘Wandering’s CC is ≥ 0.4. ‘Ranging’ is determined when it achieves the highest CC (with CC ≥ 0.7) and is described by a logistic equation (mod=32 in ‘FlexParamCurve’ package). ‘Full Round-trip’ and ‘Partial Round-trip’ are characterised by non-monotonic curves with 7 and 6 parameters, respectively, with their classification necessitating a minimum CC of ≥ 0.7. Distinction between ‘Round-trip’ and ‘Partial Round-trip’ hinges on the NSD value at the first inflection point (Morelle et al. 2017).

We also applied additional consideration for the ‘Ranging’ classification, particularly when dealing with mod2 or mod22, even if they exhibit the highest CC (Morelle et al. 2017). In such cases, these models are designated as ‘Ranging’ if the final enhanced prediction – calculated as 1.5 times the last predicted value – equals or surpasses the model’s maximum predicted value. This implies a broad movement scope, justifying the ‘Ranging’ label. It highlights situations where, despite a high CC, the movement pattern suggests a more expansive spatial reach in its concluding phase relative to the rest of the model’s predictions.

In the BCPA analysis, selecting the appropriate cluster width influences the segmentation into distinct phases because it balances capturing meaningful biological patterns against the risk of generating too many phases with little to no biological relevance (Gurarie 2014). Conversely, excessively broadening this parameter might result in a loss of critical information, as distinct movement patterns could be merged inappropriately. We identified the ideal cluster width by aligning it with the average break duration across all animals, aiming for a mean duration of approximately 120 hours. This specific duration was chosen because it exceeds the 48-72 hours typically needed by this subspecies to consume their medium-sized prey (Farhadinia et al. 2018). Such a timeframe helps prevent the misclassification of inactive periods post-hunting as relevant behavioural patterns. To isolate the widths of interest, we explored values from 15 to 70, in 5 units increments. Subsequently, to conduct the analysis over multiple cluster widths, we employed a loop that applied the classification to various cluster widths for each animal, subsequently selecting a spectrum of cluster widths that encompassed all observed durations. This approach allowed us to narrow our analysis to only those cluster widths that were pertinent to our study, thus refining our classification methodology to mirror the animals’ behavioural patterns more accurately.

#### Movement metrics and attributes

We then employed a suite of metrics to analyse the movement patterns captured in the XY coordinates, using the ‘bcpa’ package. For each positional fix, we calculated three key intrinsic movement metrics: persistence velocity, turning velocity, and speed (m/s). We also compiled three spatiotemporal attributes for each mode: total distance walked, the NSD, and the mode’s duration (hours).

We then calculated the proportion of duration, total distance walked, and NSD attributed to each mode relative to the aggregate metrics across all modes (excluding those unidentified). We also aggregated consecutive trips exhibiting the same movement mode into a single, more comprehensive movement pattern. To acknowledge the continuity of certain behaviours over time, contiguous segments that were classified under the same mode were merged, with the previously outlined metrics recalculated for these combined segments.

#### Statistical analysis

We used the proportions of time allocated to each movement mode by each individual, discounting the small amounts of time that could not be allocated to any of the modes (Table S2). We used Dirichlet regression (Douma and Weedon 2019) in ‘DirichletReg’ package (Maier 2014) to test the differences in the time proportions between the wild and rewilded leopards.

For both intrinsic movement metrics and spatiotemporal attributes, we first used Linear Discriminant Analysis (LDA) using the ‘MASS’ package (Ripley et al. 2013) to explore the variability in these metrics between wild and rewilded leopards. The inputs to the analysis were first the average of each metric for each individual across all modes. We also explored whether the patterns were consistent across movement modes (applying the LDA to attributes of single modes). The effect of origin was tested using MANOVA. The influence of input metrics on the discriminant function was tested using simple Pearson’s correlation coefficients.

We then explored the variation of per-segment average intrinsic movement metrics (persistence velocity, turning velocity, and speed) and average spatiotemporal attributes (NSD, total distance walked and duration) by focussing on whether any origin differences were consistent across modes, using generalised linear mixed models (GLMM) in the ‘lme4’ package (Bates et al. 2015). Individual identity was treated as a random effect. For each metric or attribute of interest, we ran GLMM models using the origin and movement mode as predictors and included an interaction term to test if among mode differences were similar for wild and rewilded leopards. We investigated the significance of each predictor using the function ‘drop1’ with a Chi-square likelihood ratio test. In addition, the effect of sex was tested using just rewilded leopards, and we confirmed that the overall origin effects were not affected by sex by repeating analysis without females. Model marginal means were visualized with the ‘emmeans’ package (Lenth 2022), which was also used to compare the response values among pairs of modes, adjusting for multiple testing.

## RESULTS

Overall, we removed approximately the 15% of fixes due to unsuccessful GPS retrieval, with varying percentages across animals (range 0 – 34%; Table S1 & S2), leaving 66,728 GPS fixes for 20 individual leopards. After removing the first 4 days of sampling (wild animals) and checking for erroneous fixes and outliers (Table S2), we retained on average 98% of fixes across all leopards (n=65,823 GPS fixes; Tables S1 and S2).

### Path segmentation to classify movement modes

The number of segments classified as Encamped mode (84; 22.1% and 78; 32.2%, respectively) and Wandering mode (53; 13.9% and 45; 18.6%, respectively) was similar for rewilded and wild leopards. More ranging modes were detected for rewilded leopards (153; 40.2% versus 69; 28.5%). In addition, more Round-Trip modes (68; 17.9% versus 47; 19.4%), and partial Round-Trip modes (22 versus 3) were categorized for rewilded leopards comparing to wild, respectively (Figure S3 & S4; Table 2; see Supplementary Information for extended Results).

**Table 2.** Summary statistics of the duration (hours), total distance walked (meters), net squared displacement (NSD; Km) and the number of breaks (N) for each movement mode. Ratios refer to the proportion of a metric for a given movement mode out of the total for each origin based on the aggregated movement modes (only successfully classified modes considered).

| *Mode* | *Origin* | *N* | **Duration** |  |  |  | **Distance** |  |  |  | **NSD** |  |  |  |
| --- | --- | --- | --- | --- | --- | --- | --- | --- | --- | --- | --- | --- | --- | --- |
|  |  |  | *sum* | *mean* | *sd* | *Ratio* | *sum* | *mean* | *sd* | *Ratio* | *sum* | *mean* | *sd* | *Ratio* |
| Encamped | Wild | 78 | 15210.00 | 195.00 | 145.35 | 0.39 | 5421765.42 | 69509.81 | 61382.73 | 0.40 | 1613.38 | 20.68 | 35.16 | 0.18 |
| Encamped | Rewilded | 84 | 12251.00 | 145.85 | 90.94 | 0.16 | 2313089.67 | 27536.78 | 29930.32 | 0.15 | 1081.04 | 12.87 | 27.76 | 0.01 |
| Partial round-trip | Wild | 3 | 418.00 | 139.33 | 35.80 | 0.01 | 114719.29 | 38239.76 | 14175.77 | 0.01 | 48.83 | 16.28 | 12.44 | 0.01 |
| Partial round-trip | Rewilded | 22 | 2541.00 | 115.50 | 44.64 | 0.03 | 533319.48 | 24241.79 | 15406.45 | 0.03 | 1614.33 | 73.38 | 116.26 | 0.02 |
| Ranging | Wild | 69 | 11379.00 | 164.91 | 104.36 | 0.29 | 3952453.66 | 57281.94 | 47595.86 | 0.29 | 5218.49 | 75.63 | 142.32 | 0.60 |
| Ranging | Rewilded | 153 | 43285.00 | 282.91 | 238.62 | 0.57 | 8885053.44 | 58072.24 | 61639.01 | 0.57 | 72423.59 | 473.36 | 1220.74 | 0.82 |
| Round-trip | Wild | 47 | 6475.00 | 137.77 | 65.18 | 0.16 | 2172852.34 | 46230.90 | 30977.50 | 0.16 | 651.56 | 13.86 | 19.49 | 0.07 |
| Round-trip | Rewilded | 68 | 10583.00 | 155.63 | 98.17 | 0.14 | 2345838.98 | 34497.63 | 25913.04 | 0.15 | 2764.58 | 40.66 | 96.86 | 0.03 |
| Wandering | Wild | 45 | 5792.00 | 128.71 | 50.50 | 0.15 | 1915136.43 | 42558.59 | 26946.22 | 0.14 | 1237.65 | 27.50 | 43.37 | 0.14 |
| Wandering | Rewilded | 53 | 7103.00 | 134.02 | 75.27 | 0.09 | 1475092.76 | 27831.94 | 22998.26 | 0.09 | 10636.31 | 200.69 | 562.57 | 0.12 |

In rewilded leopards, the ranging mode represented 57% of total duration compared with only 16% in Encamped mode (Table 2). In contrast, the Encamped mode accounted for 39% of the total duration across all modes and animals in wild leopards, similar to Ranging mode (29%). Most of the distance walked by wild leopards (40%) was found in Encamped modes, followed by Ranging (29%), while Ranging mode in rewilded leopards represented 57% of all distance walked followed in equal proportions by Encamped and Round-Trip (15%). Ranging also accounted for the 82% of NSD in rewilded compared to a 60% in wild leopards. The Encamped mode accounted for the 18% of NSD in wild individuals and 1% in rewilded.

Average duration of Ranging mode segments was higher in the rewilded leopards, while Encamped segments tended to be shorter - similar average duration was observed for the Wandering mode segments (Table 2). The observed pattern in average proportions across individuals reflected these numbers – the rewilded leopards tended to spend more time ranging (mean=48.6% ± SE 6.2) than did the wild leopards (mean=29.9% ± SE 5.3), In contrast, the opposite pattern was seen for the amount of time spent Encamped (mean=18.8% ± SE 3.2 vs. 38.2% ± SE 5.6) for rewilded and wild leopards, respectively. Rewilded leopards also tended to spend more time in Round-trip and Partial round-trip modes (Figure 2). The Dirichlet regression demonstrated strong evidence for country differences in mean proportions for all modes (|z| ≥ 3.4, P < 0.001), except for Partial round-trip (P=0.29).

**Figure 2.**
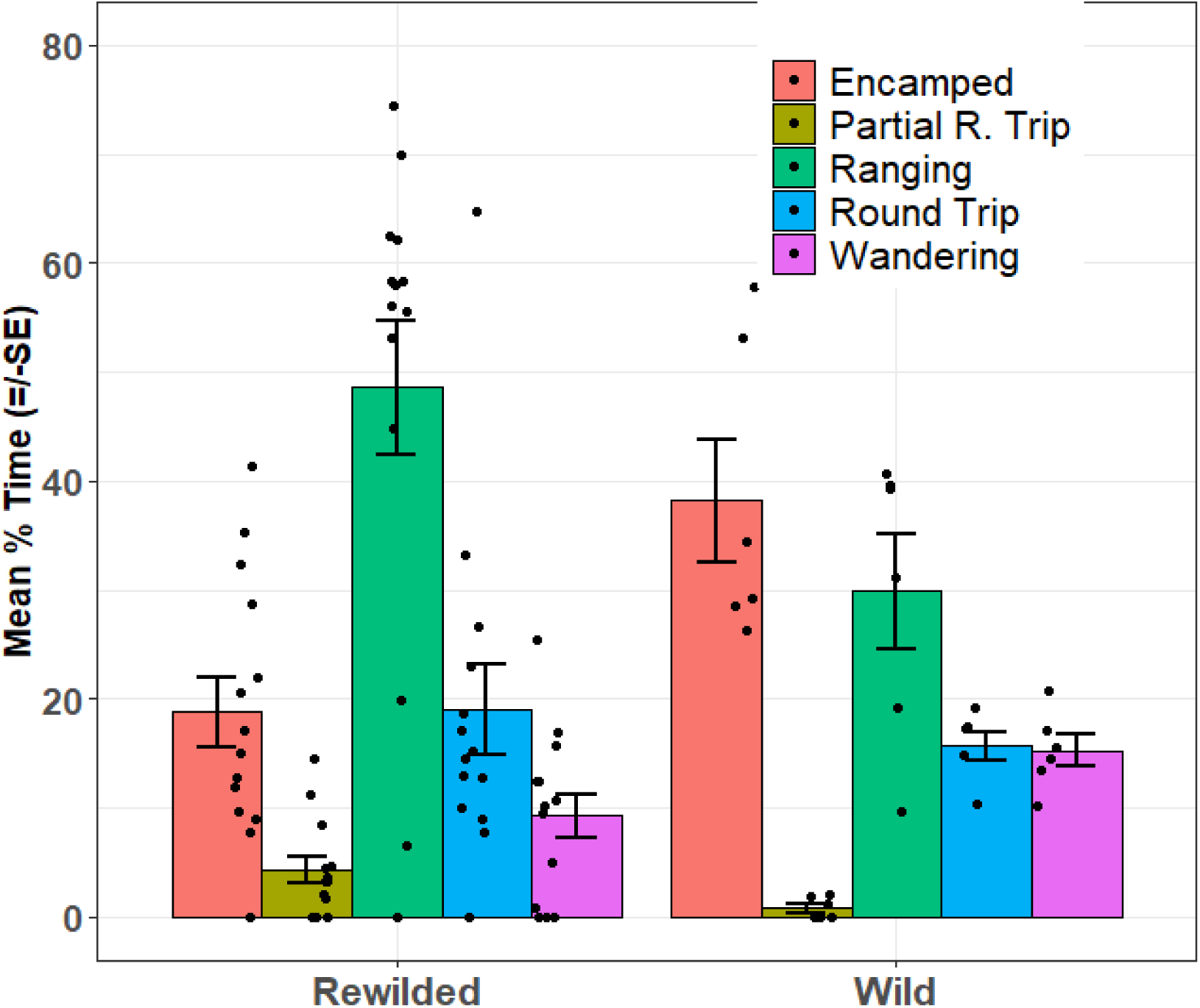
Mean % time (+/- SE) for rewilded and wild leopards allocated to the different movement modes. Points represent individuals.

### Movement metrics and attributes

The LDA demonstrated strong evidence for separation by the origin using intrinsic movement metrics (Wilk’s lambda=0.44, df=3,4, P=0.004; Figure 3A). Mean speed was the most influential variable (correlation with discriminant function r=-0.87, P<0.001). Mean persistence velocity was also negatively correlated with the discriminant function (r=-0.68, P<0.001), while the effect of turning velocity was not significant (r=0.09, P>0.05).

**Figure 3.**
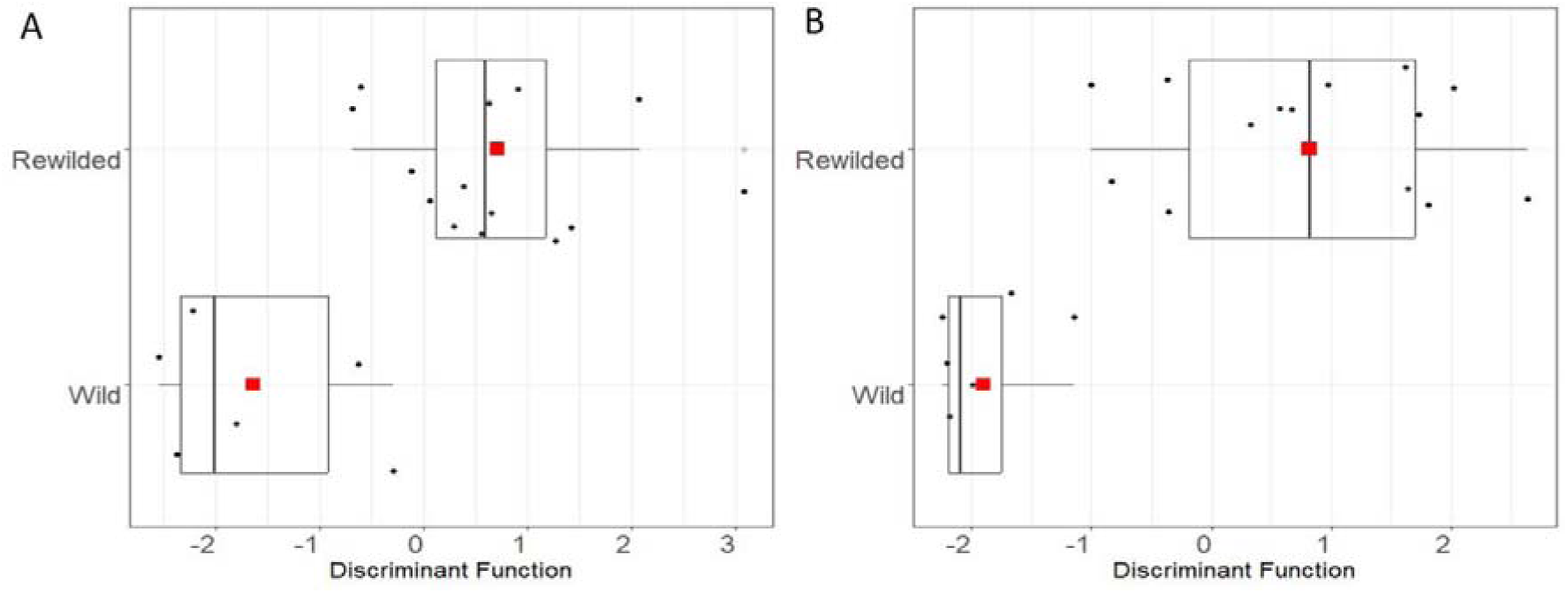
A) Discriminant function analysis using A) three intrinsic attributes of movement - mean speed, mean persistence velocity and mean turning velocity and B) three spatiotemporal attributes of movement – total step length, duration, and NSD (red points=means).

In the GLMM exploring the speed effect, there was some evidence for an interaction between the origin and mode (X^2^=8.4, df=4, P=0.07) suggestive of a different inter-mode pattern based on the individual origin. There was also strong evidence for a main effect of the origin (X^2^=7.6, df=1, P=0.005), with significantly lower speeds for the rewilded leopards. Separate analyses by the origin provided no evidence for differences between modes for wild leopards (X^2^=1.5, df=4, P=0.81), while there was evidence that speed differed between modes for rewilded leopards (X^2^=16.1, df=4, P=0.003; Figure 4A), with speeds in Encamped mode being lower than in other modes.

**Figure 4.**
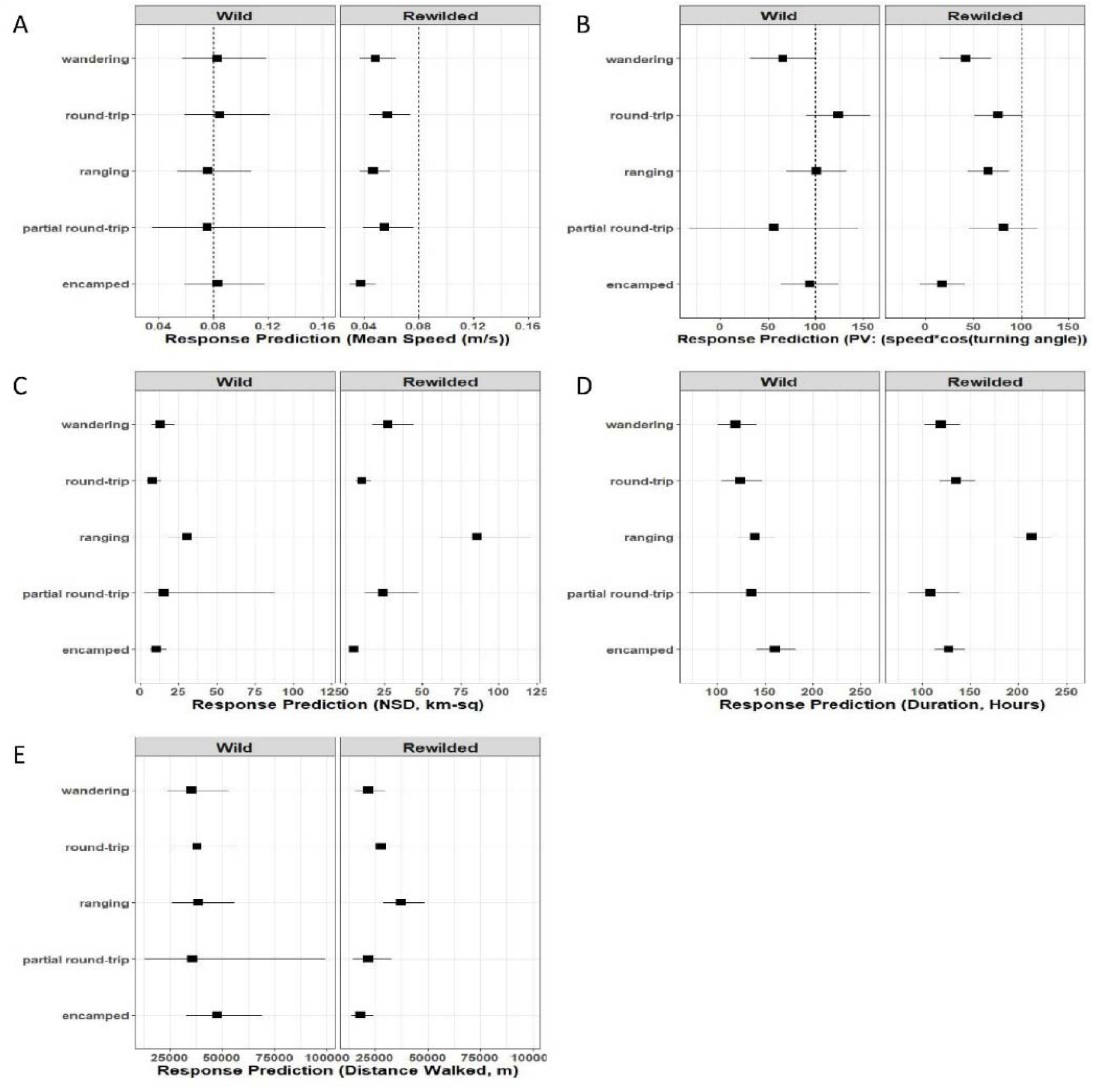
A) Estimated model marginal means for origin*mode effect in mixed model modelling mean speed. Dotted reference line at approximate mean for wild (Iranian leopards). Analysis modelled log-transformed response, back-transformed here. B) Estimated model marginal means for origin*mode effect in mixed model modelling persistence velocity. Dotted line at reference point of 100.0 added for ease of comparison. C) Estimated model marginal means for origin*mode effect in mixed model modelling NSD (in km). Log-transformed values modelled, back transformed here. D) Estimated model marginal means for origin*mode effect in mixed model modelling segment duration. Log-transformed values modelled, back transformed here. E) Estimated model marginal means for origin*mode effect in mixed model modelling total distance walked. Log-transformed values modelled, back transformed here.

Turning velocity did not appear to be associated with either origin or movement mode. There was weak evidence for an interaction between origin and movement mode (X^2^=8.0, df= 4, P= 0.09). Based on the main effects model, both effects were non-significant (X^2^=0.04, df=1, P=0.86 and X^2^=5.1, df=4, P=0.28 respectively). Persistence velocity was correlated with mean speed (r=0.35, P < 0.001), and patterns showed some similarity. There was evidence for an interaction between origin and type of mode (X^2^=11.99, df=4, P=0.02), and strong evidence for main effects of both origin and type of mode (X^2^=35.3, df=4, P < 0.001 and X^2^=7.3, df=4, P=0.007 respectively). Rewilded leopards clearly had lower overall persistence velocity (Figure 4B), particularly during Encamped and Ranging modes. Inspection of pairwise means for modes in each group of leopards separately showed persistence velocity in Encamped mode to be significantly lower than all other modes except Wandering for rewilded leopards, while no such differences are apparent in wild leopards.

### Spatiotemporal attributes

The LDA demonstrated strong evidence for separation by origin using the three spatiotemporal attributes (Wilk’s lambda=0.36, df=3,4, P < 0.001; Figure 3B). Total duration was the most influential variable (correlation with discriminant function=0.62, P < 0.001). NSD was also positively correlated with the discriminant function (r=0.55, P < 0.001), while the correlation with total distance walked was negative (r=-0.42, P < 0.001).

The models exploring individual attributes revealed some distinct effects of movement mode. The patterns were similar across the three attributes, as these attributes were correlated (|r| ≥ 0.45, P < 0.001). For NSD, there was strong evidence for an interaction between origin and mode (X^2^=31.5, df=4, P < 0.001), and no evidence for an overall effect of origin (X^2^=2.7, df=1, P=0.15). Values of NSD in ranging mode were much higher for the rewilded leopards (Figure 4C). For the wild leopards, the only significant pairwise comparisons were those including Ranging mode values, where these were significantly higher than those for encamped, Round-trip and Wandering. For the rewilded leopards, all comparisons with Ranging were significant, and encamped values were also significantly lower than those for Wandering and Partial round-trip.

For segment duration, while there was some evidence that values for the rewilded leopards were generally higher (X^2^=3.2, df=1, P=0.07), there was again strong evidence for an interaction term between origin and mode type such that the origin difference varied across mode (X^2^=30.4, df=1, P < 0.001). For the wild leopards, the only significant pairwise comparisons between modes of movements are that Encamped durations are significantly higher than Wandering. For the rewilded leopards, Ranging durations were significantly higher than for all other modes. The origin effect is therefore most marked for duration of

Ranging segments which are much higher than those for wild leopards (Figure 4D). Similarly, there was some evidence for an overall difference between the two groups, with total distance walked in rewilded leopards tending to be shorter (X^2^=4.5, df=1, P=0.03), but with strong evidence for an origin*mode interaction effect (X^2^=26.4, df=4, P < 0.001). In contrast to the patterns for NSD and duration, the distances walked were similar across modes for wild leopards but varied between modes for the rewilded leopards. For rewilded leopards total distance walked was longer in encamped modes compared with ranging and Round-trip modes, and longer in Ranging than Wandering. All the pairwise comparisons were non-significant for the wild leopards (Figure 4E).

There was no evidence for any sex effects (tested with rewilded leopards only) with the exception that rewilded female leopards had significantly longer segment durations than males (Male: 183.0 ± SE 11.1 versus female: 217 ± SE 14.6 hours; X^2^=6.1, P=0.013). There was no evidence that this differed across modes (X^2^=4.5, df=4, P=0.56).

## DISCUSSION

Our study revealed distinct behavioural and movement differences between rewilded and wild leopards. Rewilded leopards exhibited both exploratory and cautious behavioural strategies, ranging more widely over extended periods. This behaviour can be plausibly explained by the hypothesis that they were gathering information from their environments. These activities occurred at lower speeds, presumably to avoid potential risks, suggesting that they are in the process of adapting to new environments. In contrast, wild leopards demonstrated more stable and consistent movements, reflecting their familiarity with native habitats.

These differences are linked to variations in two main types of factors: landscape conditions and the individual’s features (age and origin), with the latter representing prior knowledge. We expect that the behavioural differences are partially influenced by landscape-specific ecological factors related to the distinct environments in which they reside. However, given the high ecological similarities between the two study systems, individual origin, and possibly age, should be a significant factor in explaining the differences between wild and rewilded leopards.

### Exploratory strategy hypothesis

In line with our first hypothesis, rewilded leopards spent more time in the Ranging mode. They also exhibited lower persistence velocities, especially during Encamped and Ranging modes, indicating overall slower movement with less direct and more exploratory movement paths, a pattern also observed in younger wild leopards (Farhadinia et al. 2020). Similarly, rewilded leopards dedicated more time to Round-trip and Partial Round-trip modes (Rozhnov et al. 2024). This confirms our expectation that exploratory behaviour would be conspicuous, which is consistent with studies of other reintroduced felids, such as cheetahs (*Acinonyx jubatus*) (Sievert et al. 2022), tigers (*P. tigris*) (Rozhnov et al. 2021) and lions (*P. leo*) (Hunter 1998, Kilian 2003). In contrast, resident wild leopards spent more time in the Encamped mode, indicating lower mobility, perhaps due to their greater spatial knowledge, allowing for more efficient use of resource-rich areas. This pattern can be interpreted in terms of a distinction between updating an existing cognitive map and building a new one, a fundamental aspect of animal spatial learning (Fagan et al. 2013).

The rewilded leopards also exhibited lower persistence velocities, especially during Encamped and Ranging modes. Therefore, they show slower overall movement and indicates less direct and more exploratory movement paths, a pattern also observed in non-resident wild leopards (Farhadinia et al. 2020). Nonetheless, the rewilded leopards exhibited slower movement as they ranged more widely over extended periods, plausibly explained if they were gathering essential information from their environments while cautiously avoiding potential risks. Slower movement also allowed the rewilded leopards to offset the higher energy demands of extended ranging. Movement speed also depends on habitat structure and resource dispersion. Similarly, in reintroduced lions, cumulative home ranges continued to increase over time as individuals expanded their range of exploration, albeit with different movement strategies employed by different prides (Yiu et al. 2015). Similar spatial use patterns were observed in rewilded tigers, as they exhibited expanding or shifting home ranges (Rozhnov et al. 2021) and preyed regularly on smaller species compared to wild tigers but demonstrated a higher kill rate (Miquelle et al. 2025).

### Cautious strategy hypothesis

Aligned with our second hypothesis, rewilded leopards moved at lower speeds in directional movement with less persistence in movement metrics. They also showed more frequent and larger displacements, spending more time, particularly in the Ranging mode, compared to wild leopards, reflecting more extensive and prolonged movements. These behaviours could plausibly be interpreted as indicating more cautious behaviour, which is associated with minimising risk when exploring the new environment, particularly given that many rewilded leopards wandered near human settlements (Hernandez-Blanco et al. 2024). To ensure leopards exhibit risk avoidance behaviour towards humans, they were assessed for their avoidance of humans and livestock before release. Only those that successfully passed these tests were deemed suitable for reintroduction (Rozhnov et al. 2020, 2022). Females exhibited longer segment durations than did males, suggesting sex-based differences in movement patterns in rewilded leopards. This cautious approach is frequently associated with “shy” personality trait (Spiegel et al. 2017) and may improve survival during the critical early stages of reintroduction. This pattern of cautious movement in translocated individuals has been documented across various felids species such as reintroduced lions (Yiu et al. 2015) and captive-born Iberian lynx (Cisneros-Araujo et al., 2024).

In contrast, wild leopards exhibited faster and more directed movements, reflecting their established familiarity with their environment and the need to uphold territorial boundaries. They also maintained relatively consistent speeds and showed less variability in their intrinsic movement metrics across different modes. Importantly, their displacement and segment duration demonstrated less variation, particularly during the Ranging mode, where wild leopards had smaller displacements compared to rewilded individuals. These observations suggest familiarity with their environment and an advanced spatial memory.

### Confounding factors

We analysed our dataset based on the origins of individuals, distinguishing between wild and rewilded leopards. Potential confounding factors related to distinctive landscapes may contribute to the observed behavioural differences. However, all study areas are montane rocky landscapes. Unlike the Iranian steppe site, the Russian study areas are predominantly temperate forests at lower altitudes with alpine habitats in the highlands (Rozhnov et al., 2022 2020, 2024). Similarly, medium-sized wild ungulates constitute most of the biomass consumed by leopards in all study areas (Farhadinia et al. 2018, Hernandez-Blanco et al. 2024). Nonetheless, large-sized wild ungulates such as red deer (*Cervus elaphus*) contributed to only 5% of the prey items killed by rewilded leopards in Russia (Hernandez-Blanco et al. 2024).

Additionally, rewilded leopards in Russian sites share their habitat with other large predators such as brown bears and grey wolves, while wolves are occasional in Tandoureh, Iran (Farhadinia et al. 2017a, Hernandez-Blanco et al. 2024). Although these predators can affect leopards through interference competition over kills (Hernandez-Blanco et al. 2024), it is unlikely to expect exploitation competition over leopards, as they primarily inhabit different habitat types. We therefore assumed that leopards’ mobility was not mediated by the presence of sympatric carnivores. Given the high landscape similarities between the study areas, we expect that the effect of landscape on observed behavioural differences is minor.

Rewilded leopards were released into areas without conspecifics (Rozhnov et al. 2022), whereas Iran’s study site hosts an existing leopard population (Farhadinia et al. 2019). The lack of signals from conspecifics may serve as a trigger, prompting exploratory behaviour to locate them (Naumov 1973). The presence of conspecifics can lead to extensive exploratory movements in leopards (Weise et al. 2015, Briers-Louw et al. 2019, Power et al. 2021). However, in our rewilded sample, these behaviours were observed despite the absence of a resident leopard population in Russia.

Age could also be a confounding factor in the observed behavioural differences. Although all individuals in this study were adults, those from the rewilded origin were generally younger than the wild ones. Translocation of adult individuals is typically associated with fast settlement within the first few months in felids (Sarkar et al. 2016, Dunston et al. 2017, Topličanec et al. 2022). Young wild leopards also show slower, less direct, and more exploratory movement paths (Farhadinia et al. 2020, Power et al. 2021), highlighting the confounding effects of age and origin. Given that younger animals are generally recommended for felid translocations (Power et al. 2021, Thomas et al. 2023), we suggest including a wide range of ages in translocation efforts to carefully evaluate the interaction between age and origin.

## Conclusion

Felid translocations have been practiced for decades to mitigate human-wildlife conflict (Ruth et al. 1998, Weise et al. 2015), rehabilitate injured/orphaned individuals (Thomas et al. 2023), and reinforce fragile or recovering populations (Cisneros-Araujo et al., 2024). Our recommendations for better management include:

1. Our results, combined with evidence from various felid reintroductions (Ruth et al. 1998, Houser et al. 2011, Briers-Louw et al. 2019, Cisneros-Araujo et al. 2024), confirm that translocating young dispersing individuals is associated with promising outcomes in terms of surviving in the new environment.
2. Felid reintroduction necessitates flexible, adaptive management approaches, with gradual release strategies playing a key role. These strategies enable rewilded individuals to establish stationary home ranges, develop spatial learning, and contribute to the formation of a biological signalling field at the release site (Naumov 1973). The varied responses of released animals to new environments highlight the importance of the biological signalling field in shaping population spatial organisation. Additionally, the development of a conspecific communication network helps anchor newly released individuals to the area. However, these animals do not respond uniformly to new environments. For example, in Namibian low-density leopard areas, translocated leopards quickly established home ranges (Weise et al. 2015), while in South Africa’s denser leopard populations, released individuals exhibited delayed spatial fidelity (Weilenmann et al. 2010). In Malawi, some reintroduced leopards demonstrated quick home range establishment in the absence of conspecifics, with site fidelity within 4-8 months (Briers-Louw et al., 2019). In the Russian study sites, in the absence of the biological signalling field, leopards showed no tendency to establish home ranges within the first year after release. This highlights the need for flexible management and the anticipation of dynamic behavioural plasticity in animals introduced to new environments.
3. Adopting both exploratory and cautious behavioural strategies by rewilded leopards underscores the importance of selecting low-risk areas to support the crucial stage of spatial memory development, especially for reintroducing captive-born individuals.
4. Leopards remain more exploratory in their new environment until the end of the first year after reintroduction, exhibiting persistent Ranging mode. This extended adaptation period aligns with findings from multiple felid studies, which suggest that conventional short-term monitoring periods may be insufficient to observe spatial fidelity, lifetime survival, and reproduction. For instance, breeding events—one of the indicators of successful establishment—often occur two years post-release (Houser et al. 2011, Briers-Louw et al. 2019).

## ACKNOWLEDGMENTS

We thank the Iranian Department of Environment for administrative support. We are grateful to the experts at the Sochi Breeding Center, Russia, particularly U.A. Semenov. and M.V.Alshinetskiy for veterinarian assistance. S.G. Shevelev, M.M. Gatsiev and S.K. Dzgoev who supported our work in the Caucasian Nature Reserve, National Park Alanya and Turmonskiy regional natural protected area. We are grateful to the field crew, particularly P. Moghadas, I. Memarian and J. Kaandorp, A. Shahrdari, and rangers for their field assistance in Iran as well as I. P. Voschanova, and Z. Dzutsev in Russia. We thank M.R.D. Magomedov (DSC RAS) for the data from Dagestan and Ch.M. Mamiev, A.B. Alibekov and M.E. Slanova for contributing to this project in Central Caucasus.

## AUTHOR CONTRIBUTION

MSF, LA, VVR, JAHB, AY and DWM conceived the ideas and designed methodology; MSF, DWM, VVR, NAD acquired funding, MSF, JAHB, AY, VVR, KH, MDC, MA, DN, AP and PW collected the data; MSF, LA and PJJ analysed the data; MSF, LA and JAHB led the writing of the manuscript. All authors contributed critically to the drafts and gave final approval for publication.

## REFERENCES

Bates, D., M. Mächler, B. Bolker, and S. Walker. 2015. Fitting linear mixed-effects models using lme4. 67 (1), 48. J Stat Softw https://doi.org/10 18637.

Bjørneraas, K., B. Van Moorter, C. M. Rolandsen, and I. Herfindal. 2010. Screening global positioning system location data for errors using animal movement characteristics. The Journal of Wildlife Management 74:1361–1366.

Börger, L., B. D. Dalziel, and J. M. Fryxell. 2008. Are there general mechanisms of animal home range behaviour? A review and prospects for future research. Ecology letters 11:637–650.

Börger, L., and J. Fryxell. 2012. Quantifying individual differences in dispersal using net squared displacement. Dispersal ecology and evolution 30:222–230.

Briers-Louw, W. D., S. Verschueren, and A. J. Leslie. 2019. Big cats return to Majete Wildlife Reserve, Malawi: evaluating reintroduction success. African Journal of Wildlife Research 49:34–50.

Bunnefeld, N., L. Börger, B. van Moorter, C. M. Rolandsen, H. Dettki, E. J. Solberg, and G. Ericsson. 2011. A modelLdriven approach to quantify migration patterns: individual, regional and yearly differences. Journal of Animal Ecology 80:466–476.

Calabrese, J. M., C. H. Fleming, and E. Gurarie. 2016. ctmm: an R package for analyzing animal relocation data as a continuousLtime stochastic process. Methods in Ecology and Evolution 7:1124–1132.

Calenge, C. 2006. The package “adehabitat” for the R software: a tool for the analysis of space and habitat use by animals. Ecological modelling 197:516–519.

Cisneros-Araujo, P., G. Garrote, A. Corradini, M. S. Farhadinia, B. Robira, G. López, L. Fernández, M. López-Parra, M. García-Tardío, and R. Arenas-Rojas. 2024. Born to be wild: Captive-born and wild Iberian lynx (Lynx pardinus) reveal space-use similarities when reintroduced for species conservation concerns. Biological Conservation 294:110646.

Cooke, S. J., J. R. Bennett, and H. P. Jones. 2019. We have a long way to go if we want to realize the promise of the “Decade on Ecosystem Restoration.” Conservation Science and Practice 1:e129.

Douma, J. C., and J. T. Weedon. 2019. Analysing continuous proportions in ecology and evolution: A practical introduction to beta and Dirichlet regression. Methods in Ecology and Evolution 10:1412–1430.

Dunston, E. J., J. Abell, R. E. Doyle, D. Duffy, C. Poynter, J. Kirk, V. B. Hilley, A. Forsyth, E. Jenkins, and D. Mcallister. 2017. Does captivity influence territorial and hunting behaviour? Assessment for an ex situ reintroduction program of African lions Panthera leo. Mammal Review 47:254–260.

Edelhoff, H., J. Signer, and N. Balkenhol. 2016. Path segmentation for beginners: an overview of current methods for detecting changes in animal movement patterns. Movement ecology 4:1–21.

Fagan, W. F., M. A. Lewis, M. AugerLMéthé, T. Avgar, S. Benhamou, G. Breed, L. LaDage, U. E. Schlägel, W. Tang, and Y. P. Papastamatiou. 2013. Spatial memory and animal movement. Ecology letters 16:1316–1329.

Falcón-Cortés, A., D. Boyer, E. Merrill, J. L. Frair, and J. M. Morales. 2021. Hierarchical, memory-based movement models for translocated elk (cervus canadensis). Frontiers in Ecology and Evolution 9:702925.

Farhadinia, M. S., P. J. Johnson, L. T. B. Hunter, and D. W. Macdonald. 2017a. Wolves can suppress goodwill for leopards: Patterns of human-predator coexistence in northeastern Iran. Biological Conservation 213.

Farhadinia, M. S., P. J. Johnson, L. T. B. Hunter, and D. W. Macdonald. 2018. Persian leopard predation patterns and kill rates in the Iran–Turkmenistan borderland. Journal of Mammalogy 99:713–723.

Farhadinia, M. S., B. T. McClintock, P. J. Johnson, P. Behnoud, K. Hobeali, P. Moghadas, L. T. B. Hunter, and D. W. Macdonald. 2019. A paradox of local abundance amidst regional rarity: the value of montane refugia for Persian leopard conservation. Scientific Reports 9.

Farhadinia, M. S., I. Memarian, K. Hobeali, A. Shahrdari, B. Ekrami, J. Kaandorp, and D. W. Macdonald. 2017b. GPS collars reveal transboundary movements by Persian leopards in Iran. Cat News 65:28–30.

Farhadinia, M. S., T. Michelot, P. J. Johnson, L. T. B. Hunter, and D. W. MacDonald. 2020. Understanding decision making in a food-caching predator using hidden Markov models. Movement Ecology 8.

Gigliotti, L. C., M. R. Matchett, and D. S. Jachowski. 2019. Mountain lions on the prairie: habitat selection by recolonizing mountain lions at the edge of their range. Restoration Ecology 27:1032–1040.

Gupte, P. R., C. E. Beardsworth, O. Spiegel, E. Lourie, S. Toledo, R. Nathan, and A. I. Bijleveld. 2022. A guide to preLprocessing highLthroughput animal tracking data. Journal of Animal Ecology 91:287–307.

Gurarie, E. 2014. bcpa: Behavioral change point analysis of animal movement. R package version 1.1.

Gurarie, E., R. D. Andrews, and K. L. Laidre. 2009. A novel method for identifying behavioural changes in animal movement data. Ecology letters 12:395–408.

Gurarie, E., C. Bracis, M. Delgado, T. D. Meckley, I. Kojola, and C. M. Wagner. 2016. What is the animal doing? Tools for exploring behavioural structure in animal movements. Journal of Animal Ecology 85:69–84

Hernandez-Blanco, J. A., M. D. Chistopolova, A. B. Pkhitikov, S. A. Trepet, P. I. Weinberg, Z. V Dzutsev, A. A. Yachmennikova, N. A. Dronova, A. N. Minaev, and S. V Naidenko. 2024. A comprehensive analysis of the hunting behavior of reintroduced leopards (Panthera pardus ciscaucasica) in the Russian Caucasus. Biology Bulletin 51:S342–S353.

Houser, A., L. K. Boast, M. J. Somers, M. Gusset, and C. J. Bragg. 2011. Pre-release hunting training and post-release monitoring are key components in the rehabilitation of orphaned large felids. South African Journal of Wildlife Research-24-month delayed open access 41:11–20.

Hunter, L. T. B. 1998. Early post-release movements and behaviour of reintroduced cheetahs and lions, and technical considerations in large carnivore restoration. Page Proceedings of a Symposium on Cheetahs as game ranch animals.

IUCN. 2013. Guidelines for reintroductions and other conservation translocations. Page Gland Switzerland Camb UK IUCNSSC Re-Introd Spec Group. Gland, Switzerland / Cambridge, UK.

Kilian, P. J. 2003. The ecology of reintroduced lions on the Welgevonden Private Game Reserve, Waterberg. University of Pretoria (South Africa).

Komsta, L., and F. Novomestky. 2015. Moments, cumulants, skewness, kurtosis and related tests. R package version 14.

Lenth, R. 2022. emmeans: Estimated marginal means, aka least-squares means. R package version 1.7. 2.

Maier, M. 2014. DirichletReg: Dirichlet regression for compositional data in R.

Miquelle, D. G., A. S. Mukhacheva, E. V Bragina, S. J. Waller, Y. K. Petrunenko, S. V Naidenko, J. A. HernandezLBlanco, V. A. Kastrikin, A. N. Rybin, and N. N. Rybin. 2025. Rehabilitating tigers for range expansion: lessons from the Russian Far East. The Journal of Wildlife Management 89:e22691.

Van Moorter, B., D. Visscher, S. Benhamou, L. Börger, M. S. Boyce, and J. Gaillard. 2009. Memory keeps you at home: a mechanistic model for home range emergence. Oikos 118:641–652.

Morelle, K., N. Bunnefeld, P. Lejeune, and S. A. Oswald. 2017. From animal tracks to fineLscale movement modes: a straightforward approach for identifying multiple spatial movement patterns. Methods in Ecology and Evolution 8:1488–1498.

Naumov, N. P. 1973. Signal (biological) fields and their significance for animals. Zhurnal obshchei biologii 34:808–817.

Oswald, S. A., I. C. T. Nisbet, A. Chiaradia, and J. M. Arnold. 2012. FlexParamCurve: R package for flexible fitting of nonlinear parametric curves. Methods in Ecology and Evolution 3:1073–1077.

Popov, S. V. 2010. Environmental uncertainty and arousal/stress as the direct determinants of animal behaviour. Zhurnal Obshchei Biologii 71:287–297.

Power, R. J., L. Venter, M. Botha, and P. Bartels. 2021. Repatriating leopards into novel landscapes of a South African province. Ecological Solutions and Evidence 2:e12046.

R Development Core Team. 2013. R: A language and environment for statistical computing.

Ranc, N., F. Cagnacci, and P. R. Moorcroft. 2022. Memory drives the formation of animal home ranges: evidence from a reintroduction. Ecology Letters 25:716–728.

Ripley, B., B. Venables, D. M. Bates, K. Hornik, A. Gebhardt, D. Firth, and M. B. Ripley. 2013. Package ‘mass.’ Cran r 538:113–120.

Rozhnov, V., and V. Lukarevsky. 2008. Program for reintroduction of Central Asian leopard in the Caucasus region. Moscow: KMK:65.

Rozhnov, V., S. Naidenko, J. Hernandez-Blanco, M. Chistopolova, P. Sorokin, A. Yachmennikova, E. Y. Blidchenko, A. Y. Kalinin, and V. A. Kastrikin. 2021. Restoration of the Amur tiger (Panthera tigris altaica) population in the northwest of its distribution area. Biology Bulletin 48:1401–1423.

Rozhnov, V. V, A. A. Yachmennikova, I. P. Kotlov, D. D. Arsanukaev, E. A. Aristarkhova, M.-R. Magomedov, P. I. Weinberg, J. A. Hernandez-Blanco, M. D. Chistopolova, and N. A. Dronova. 2024. Restoration of the Persian Leopard in the Russian Caucasus: Suitability Modeling of Prey Base and Competitors, Habitat Fragmentation Assessment. Biology Bulletin 51:S242–S265.

Rozhnov, V., A. Yachmennikova, N. Dronova, S. Naidenko, J. Hernandez-Blanco, M. Chistopolova, A. Pkhitikov, F. Tembotova, S. Trepet, and I. Chestin. 2022. Experience of the leopard re-covering through reintroduc-tion in the Russian Caucasus. CATnews.

Rozhnov, V., A. Yachmennikova, N. Dronova, A. Pkhitikov, M. Magomedov, I. Chestin, and Alibekov. AB. 2020. The restoration of Persian leopard in the Caucasus (scientific approach). KMK Scientific Press Ltd., Moscow.

Ruth, T. K., K. A. Logan, L. L. Sweanor, M. G. Hornocker, and L. J. Temple. 1998. Evaluating cougar translocation in New Mexico. The Journal of wildlife management:1264–1275.

Sarkar, M. S., K. Ramesh, J. A. Johnson, S. Sen, P. Nigam, S. K. Gupta, R. S. Murthy, and G. K. Saha. 2016. Movement and home range characteristics of reintroduced tiger (Panthera tigris) population in Panna Tiger Reserve, central India. European Journal of Wildlife Research 62:537–547.

Sievert, O., J. Fattebert, K. Marnewick, and A. Leslie. 2022. Assessing the success of the first cheetah reintroduction in Malawi. Oryx 56:505–513.

Singh, N. J., L. Börger, H. Dettki, N. Bunnefeld, and G. Ericsson. 2012. From migration to nomadism: movement variability in a northern ungulate across its latitudinal range. Ecological Applications 22:2007–2020.

Soleymani, A., F. Pennekamp, S. Dodge, and R. Weibel. 2017. Characterizing change points and continuous transitions in movement behaviours using wavelet decomposition. Methods in Ecology and Evolution 8:1113–1123.

Spiegel, O., S. T. Leu, C. M. Bull, and A. Sih. 2017. What’s your move? Movement as a link between personality and spatial dynamics in animal populations. Ecology letters 20:3– 18.

Sulikowski, D., and D. Burke. 2011. Movement and memory: different cognitive strategies are used to search for resources with different natural distributions. Behavioral Ecology and Sociobiology 65:621–631.

Svenning, J. C., P. B. M. Pedersen, C. J. Donlan, R. Ejrnæs, S. Faurby, M. Galetti, D. M. Hansen, B. Sandel, C. J. Sandom, J. W. Terborgh, and F. W. M. Vera. 2016. Science for a wilder Anthropocene: Synthesis and future directions for trophic rewilding research. Proceedings of the National Academy of Sciences of the United States of America 113:898–906.

Thomas, S., V. van der Merwe, W. D. Carvalho, C. H. Adania, R. Černe, T. Gomerčić, M. Krofel, J. Thompson, R. T. McBride, J. Hernandez-Blanco, A. Yachmennikova, D. W. Macdonald, and M. S. Farhadinia. 2023. Evaluating the performance of conservation translocations in large carnivores across the world. Biological Conservation 279.

Topličanec, I., T. Gomerčić, M. Krofel, I.-M. Pop, J. Kubala, B. Tám, S. Blašković, and M. Sindičić. 2022. Early post-release behaviour of Eurasian lynx translocated to the transboundary region of the Dinaric Mountains. Journal of Vertebrate Biology 71:22061–22064.

Weilenmann, M., M. Gusset, D. R. Mills, T. Gabanapelo, and M. Schiess-Meier. 2010. Is translocation of stock-raiding leopards into a protected area with resident conspecifics an effective management tool? Wildlife Research 37:702–707.

Weise, F. J., J. Lemeris, K. J. Stratford, R. J. van Vuuren, S. J. Munro, S. J. Crawford, L. L. Marker, and A. B. Stein. 2015. A home away from home: insights from successful leopard (Panthera pardus) translocations. Biodiversity and conservation 24:1755–1774.

Wong, B. B. M., and U. Candolin. 2015. Behavioral responses to changing environments. Behavioral Ecology 26:665–673.

Yiu, S.-W., M. Keith, L. Karczmarski, and F. Parrini. 2015. Early post-release movement of reintroduced lions (Panthera leo) in Dinokeng Game Reserve, Gauteng, South Africa. European Journal of Wildlife Research 61:861–870.

