## Supplementary Information for "Cautious explorers: comparing movement patterns of wild and rewilded solitary predators"

### Extended Materials and methods

#### Leopard handling and collaring

In Iran, we captured leopards with Aldrich foot-snares extensively modified to reduce chances of injury and remotely monitored with VHF trap transmitters (Wildlife Materials, Inc., Illinois, USA) every 1–2 hours. As leopards are known to respond to baits, a wild pig carcass was used as bait, normally hanging from a tree or rock. Traps were also deployed along trails leading to the baits. In summer, we deployed traps along trails leading to water sources, sometimes without bait (see Farhadinia et al. (2017) for more details).

We immobilized leopards using a combination of ketamine 10% (Alfasan, Nederland BV) 2 mg/kg, medetomidine HCl 20 mg/ml (Kyron Laboratories (Pety) Ltd., Johannesburg, South Africa) 30 μg/kg and butorphanol 0.2 mg/kg (Torbugesic®, Fort Dodge Animal Health Fort Dodge Animal Health, Iowa 50501 USA) delivered intramuscularly with a dart gun (Daninject, Denmark) using a 1.5 ml dart. Anesthesia lasted for 44 to 60 minutes, followed by reversal using atipamazole (3 times the medetomidine dosage) and nantroxan (the doses equal to butorphanole), injected intramuscularly.

Age estimates were based on dental features (Farhadinia et al. 2014). We used GPS collars with Iridium download (LOTEK Engineering Ltd., Newmarket, ON, Canada). Each collar incorporated a drop-off buckle with a timer set to 52 weeks since deployment. Collars weighed 640 g, equivalent to less than 1–2% of leopard body mass. For programming the collars’ fix rates, we recorded fixes every 2-3 hours. However, to increase fix success rates, fixes were taken hourly during the last week of each month. Also, a ‘virtual fence’ option enabled us to upload the area’s boundary, so that when leopards left the defined area fix rate could be increased to hourly.

In Russia, all leopards were born in captivity in the Sochi Leopard Restoration Center and were released within two study sites (Table 1) at ages ranging between 20 to 36 months, following at least of training in captivity (see Rozhnov et al. (2020) for further details; see the details of released leopards in Table S1). Several weeks before the release, the leopards were immobilised to fit a collar with an intramuscular injection of 2.5 mg/kg of Zoletil (Tiletamine hydrochloride and Zolazepam hydrochloride, Virbac, France) mixed with 0.02 mg/kg Domitor (Medetomidine hydrochloride, Pfizer, USA). If necessary, Domitor was reversed by Antisedan (Atipamezole hydrochloride, Pfifer USA) with a dosage of 0.02 mg/kg. We fitted the leopards with Lotek GPS Iridium track M (three collars in 2016), Lotek GPS Litetrack Iridium (three collars in 2018 and four in 2020) (LOTEK Engineering Ltd., Newmarket, ON, Canada) and Moosefarm GPS-GSM (IEE RAS, Russia; including three and one collars in 2022 and 2023, respectively). We programmed the collars to obtain 12 (Lotek Iridiumtrack) and 24 GPS (Lotek Litetrack and Moosefarm) fixes per day. Before being transported to the release site, the leopards were also anaesthetised and placed in standard transport cages. The transport was carried out by helicopter MI-8, following the animals recovered from the anaesthesia. The leopards were released directly into the wild without an intermediate enclosure for adaptation (hard release). Details of leopards as well as their tracking periods are provided on Table S2. Out of 14 leopards released, 12 survived until 6 months post-release, and 8 were still alive by the end of their first year (Table S1; see Supplementary Information for further details).

#### Data screening

We omitted the first 4 days for all collar data from Iran, associated with the earliest known kill made by the leopards after collaring. Previous work has shown than fixes obtained immediately after collaring should be excluded because the animal is likely to behave abnormally, e.g. less mobility due to the prolonged effects of immobilization after reversal (Bjørneraas et al. 2010). GPS fixes were inspected for possible errors, including missing locations fixes (unsuccessful attempts of a GPS fix) and outliers.

Erroneous locations and outliers were inspected in a three-steps fashion. First, we adopted the script developed by Bjørneraas et al. (2010)’s aimed at the identification of unrealistic movement patterns, likely deriving from GPS errors. In summary, the method relies on a predefined distance threshold that is possible for a given animal to travel within the sampling interval, and on the identification of spikes based on the speed and turning angles between consecutive fixes. For this part, we adopted the conservative parameters defined for Persian leopards by Farhadinia et al. (2018), as Δ = 30,000 m; μ = 15,000 m; α = 5000 m/h; θ = −0.97 corresponding to turning angles between 166˚ and d 194˚; Δ is a distance threshold over which an individual could not possibly travel between consecutive intervals, μ is a distance that leopard can move between two fixes and α is speed.

Concurrently, we screened the distribution of speeds between consecutive fixes using the ‘atlastools’ package (Gupte et al. 2022), and we isolated all those fixes whose speeds and turning angles were above the 95^th^ percentile of the values distribution. After having removed erroneous fixes identified both by Bjørneraas et al.’s (2010) method and ‘atlastools’ outliers, we visually inspected all tracks using the function ‘outlie’ implemented in the ‘ctmm’ package (Calabrese et al. 2016) and removed obvious outliers without relying on distance or speed thresholds.

### Extended Results

#### Path segmentation to classify movement modes

The parameter selection process selected mainly a window size of 20 (16 animals), followed by 30 (three animals) and 40 (one animal). Step size was mostly selected at three and four, while the K parameter was mostly selected at four (Table S3).

The selection of the cluster width revealed that 30 was the value most retrieved across all leopards (13 animals), followed by 25 and 30 (three animals each) and 20 for one animal. The segmentation analyses produced a total of 986 breaks. The overall mean number of breaks (i.e. change points) per individual was 49.3 (SE=6.9) with mean duration across breaks and animals of 121.8 hours (SE=1.8). Overall, only 10 segments failed to be classified into movement modes, representing 1.0% of the whole dataset (range 0 – 9.09; Table S4 & S5).

Clumping contiguous and continuous modes into a larger spatiotemporal process reduced the number of breaks to 622. The mean number of breaks per individual then became 31.1 (SE=4.2; Table S5; Figure 2 & S3 & S4). Overall, the aggregation resulted in 162 behavioural modes classified as ‘Encamped’ (26.0%), 25 as ‘Partial Round-Trip’ (4.0%), 222 as ‘Ranging’ (35.7%), 115 as ‘Round-Trip’ (18.5%) and 98 classified as ‘Wandering’ (15.8%) (Table S6 & S7). The average duration (hours) across all animals for each movement mode was 155.2 ± 91.4 (Encamped), 85.2 ± 10.6 (Partial Round-Trip), 216.8 ± 155.8 (Ranging), 150.7 ± 74.4 (Round-Trip) and 108.2 ± 42.4 (Wandering; Table S6 & S7). Nonetheless, durations tend to be significantly shorter for wild leopards (P=0.05 and P=0.04 for original and aggregated movement modes, respectively; Table S5).

Table S1. Details on the Persian leopards monitored in this study, including age at the time of capture/release, gender, locality of capture/release, collar make and sampling lag, sampling duration, number of incomplete fixes (NA) and percentage out of the total (%NA), number of complete fixes (N), number of fixes after screening for duplicates, errors, and outliers (N filtered) and their percentage compared to the total of complete fixes (% retained). Wild leopards were monitored in Iran while rewilded leopards were studied in Russia.

| **Individual** | **Origin** | **Age** | **Sex** | **Locality** | **Collar** | **Lag** | **From** | **To** | **NA** | **%NA** | **N** | **N filtered** | **% retained** |
| --- | --- | --- | --- | --- | --- | --- | --- | --- | --- | --- | --- | --- | --- |
| **Bardia** | Wild | 8-10 years | M | Tandoureh National Park, Iran | Lotek (Iridium) | 3 | 02/10/2014 | 30/09/2015 | 370 | 0.08 | 4002 | 3917 | 0.98 |
| **Borna** | Wild | 5-6 years | M | Tandoureh National Park, Iran | Lotek (Iridium) | 1 | 27/09/2014 | 27/09/2015 | 387 | 0.08 | 4558 | 4499 | 0.99 |
| **Borzou** | Wild | >10 years | M | Tandoureh National Park, Iran | Lotek (Iridium) | 1 | 06/02/2015 | 05/02/2016 | 820 | 0.12 | 5991 | 5955 | 0.99 |
| **Iran** | Wild | 2-3 years | F | Tandoureh National Park, Iran | Lotek (Iridium) | 1 | 05/12/2015 | 29/01/2016 | 146 | 0.12 | 1030 | 1009 | 0.98 |
| **Kaveh** | Wild | 3-4 years | M | Tandoureh National Park, Iran and Turkmenistan | Lotek (Iridium) | 1 | 03/09/2015 | 26/08/2016 | 358 | 0.06 | 6106 | 6073 | 0.99 |
| **Tandoureh** | Wild | 7-10 years | M | Tandoureh National Park, Iran | Lotek (Iridium) | 1 | 15/08/2016 | 01/04/2017 | 161 | 0.05 | 2769 | 2693 | 0.97 |
| **Agura** | Rewilded | 2 years | F | Turmonskiy Wildlife Sanctuary, Russia | Lotek (Iridium) | 1 | 25/08/2020 | 22/03/2021 | 1645 | 0.33 | 3354 | 3321 | 0.99 |
| **Akhun** | Rewilded | 3 years | M | Caucasus Nature Reserve, Russia | Lotek (Iridium) | 2 | 15/07/2016 | 13/08/2016 | 42 | 0.12 | 306 | 306 | 1.00 |
| **Artek** | Rewilded | 2 years | M | Caucasus Nature Reserve, Russia | Lotek (Iridium) | 1 | 31/08/2018 | 03/01/2020 | 2645 | 0.32 | 5564 | 5495 | 0.99 |
| **Baksan** | Rewilded | 2 years | M | Turmonskiy Wildlife Sanctuary, Russia | Lotek (Iridium) | 1 | 25/08/2020 | 21/10/2020 | 448 | 0.33 | 918 | 911 | 0.99 |
| **Chilmas** | Rewilded | 2 years | M | Turmonskiy Wildlife Sanctuary, Russia | Moosefarmer (GSM) | 1 | 22/07/2022 | 27/10/2023 | 0 | 0.00 | 1215 | 1157 | 0.95 |
| **Elbrus** | Rewilded | 2 years | M | Alaniya National Park, Russia | Lotek (Iridium) | 1 | 27/07/2018 | 27/11/2018 | 759 | 0.26 | 2179 | 2151 | 0.99 |
| **Khosta** | Rewilded | 1 year 8 months | F | Turmonskiy Wildlife Sanctuary, Russia | Moosefarmer (GSM) | 1 | 16/07/2022 | 12/08/2023 | 0 | 0.00 | 5690 | 5595 | 0.98 |
| **Killi** | Rewilded | 2 years | M | Caucasus Nature Reserve, Russia | Lotek (Iridium) | 2 | 15/07/2016 | 02/09/2017 | 562 | 0.12 | 4049 | 4049 | 1.00 |
| **Kodor** | Rewilded | 2 years | M | Caucasus Nature Reserve, Russia | Lotek (Iridium) | 1 | 20/08/2020 | 01/09/2020 | 76 | 0.28 | 200 | 192 | 0.96 |
| **Laba** | Rewilded | 2 years | F | Caucasus Nature Reserve, Russia | Lotek (Iridium) | 1 | 20/08/2020 | 26/10/2020 | 299 | 0.19 | 1291 | 1265 | 0.98 |
| **Laura** | Rewilded | 1 year 8 months | F | Turmonskiy Wildlife Sanctuary, Russia | Moosefarmer (GSM) | 1 | 16/07/2022 | 24/08/2023 | 0 | 0.00 | 5306 | 5165 | 0.97 |
| **Leo** | Rewilded | 1 year 8 months | M | Turmonskiy Wildlife Sanctuary, Russia | Moosefarmer (GSM) | 1 | 16/07/2022 | 07/05/2023 | 0 | 0.00 | 4097 | 4026 | 0.98 |
| **Victoria** | Rewilded | 3 years | F | Caucasus Nature Reserve, Russia | Lotek (Iridium) | 2 | 15/07/2016 | 21/06/2017 | 418 | 0.11 | 3540 | 3538 | 1.00 |
| **Volna** | Rewilded | 2 years | F | Alaniya National Park, Russia | Lotek (Iridium) | 1 | 27/07/2018 | 02/07/2019 | 2364 | 0.34 | 4563 | 4506 | 0.99 |

Table S2. Details of the data filtering methods for the complete fixes. Columns show the number of points removed on wild captured animals (First 4 Days), Duplicate coordinates deriving from collar malfunctioning, erroneous locations respectively assessed through Bjørneraas et al. (2010)’s method, atlastools (Gupte et al. 2022), and ctmm’s outlie function (Calabrese et al. 2016).

| **Individual** | **All records** | **First 4 days** | **Duplicates** | **Bjørneraas** | **atlastools** | **ctmm** | **Final set** | **Removed** |
| --- | --- | --- | --- | --- | --- | --- | --- | --- |
| **Bardia** | 4002 | 77 | 0 | 0 | 8 | 0 | 3917 | 85 |
| **Borna** | 4558 | 48 | 0 | 0 | 11 | 0 | 4499 | 59 |
| **Borzou** | 5991 | 28 | 0 | 0 | 8 | 0 | 5955 | 36 |
| **Iran** | 1030 | 19 | 0 | 0 | 2 | 0 | 1009 | 21 |
| **Kaveh** | 6106 | 25 | 0 | 0 | 8 | 0 | 6073 | 33 |
| **Tandoureh** | 2769 | 71 | 0 | 0 | 5 | 0 | 2693 | 76 |
| **Agura** | 3354 | 0 | 0 | 5 | 28 | 0 | 3321 | 33 |
| **Akhun** | 306 | 0 | 0 | 0 | 0 | 0 | 306 | 0 |
| **Artek** | 5564 | 0 | 0 | 19 | 50 | 0 | 5495 | 69 |
| **Baksan** | 918 | 0 | 0 | 1 | 6 | 0 | 911 | 7 |
| **Chilmas** | 1215 | 0 | 58 | 0 | 0 | 0 | 1157 | 58 |
| **Elbrus** | 2179 | 0 | 0 | 8 | 20 | 0 | 2151 | 28 |
| **Khosta** | 5690 | 0 | 94 | 0 | 1 | 0 | 5595 | 95 |
| **Killi** | 4049 | 0 | 0 | 0 | 0 | 0 | 4049 | 0 |
| **Kodor** | 200 | 0 | 0 | 3 | 4 | 1 | 192 | 8 |
| **Laba** | 1291 | 0 | 0 | 11 | 14 | 1 | 1265 | 26 |
| **Laura** | 5306 | 0 | 140 | 0 | 1 | 0 | 5165 | 141 |
| **Leo** | 4097 | 0 | 71 | 0 | 0 | 0 | 4026 | 71 |
| **Victoria** | 3540 | 0 | 0 | 0 | 2 | 0 | 3538 | 2 |
| **Volna** | 4563 | 0 | 0 | 18 | 39 | 0 | 4506 | 57 |

Table S3. Parameters of BCPA selected in each leopard as the combination of settings causing less kurtosis (in either direction) in the residuals of the BCPA function.

| **Individual** | **Window** | **Step** | **K** |
| --- | --- | --- | --- |
| Bardia | 20 | 3 | 4 |
| Borna | 20 | 3 | 4 |
| Borzou | 20 | 3 | 4 |
| Iran | 20 | 4 | 4 |
| Kaveh | 20 | 4 | 4 |
| Tandoureh | 20 | 4 | 4 |
| Agura | 20 | 4 | 4 |
| Akhun | 20 | 3 | 4 |
| Artek | 30 | 3 | 2 |
| Baksan | 20 | 4 | 4 |
| Chilmas | 20 | 1 | 2 |
| Elbrus | 20 | 1 | 4 |
| Khosta | 20 | 4 | 4 |
| Killi | 30 | 3 | 2 |
| Kodor | 20 | 2 | 1 |
| Laba | 20 | 4 | 4 |
| Laura | 20 | 3 | 4 |
| Leo | 20 | 4 | 4 |
| Victoria | 30 | 4 | 4 |
| Volna | 40 | 1 | 4 |

Table S4. Number of breaks (N) found for each animal after BCPA, number of non-classified breaks (NA) and their percentage (NA%).

| **Individual** | **N** | **NA** | **NA%** |
| --- | --- | --- | --- |
| Agura | 37 | 0 | 0 |
| Akhun | 6 | 0 | 0 |
| Artek | 102 | 1 | 0.98 |
| Baksan | 11 | 0 | 0 |
| Bardia | 69 | 1 | 1.45 |
| Borna | 75 | 2 | 2.67 |
| Borzou | 71 | 0 | 0 |
| Chilmas | 19 | 0 | 0 |
| Elbrus | 27 | 0 | 0 |
| Iran | 11 | 1 | 9.09 |
| Kaveh | 76 | 2 | 2.63 |
| Khosta | 72 | 1 | 1.39 |
| Killi | 79 | 0 | 0 |
| Kodor | 2 | 0 | 0 |
| Laba | 12 | 0 | 0 |
| Laura | 80 | 0 | 0 |
| Leo | 61 | 2 | 3.28 |
| Tandoureh | 48 | 0 | 0 |
| Victoria | 64 | 0 | 0 |
| Volna | 64 | 0 | 0 |
| Total | 986 | 10 | 1.01 |

Table S5. The number of breaks and corresponding cluster width whose average duration time was the closest to the time scale of interest in this analysis (120 hours). Cluster width duration has been explored on the original interpolated dataset. The table also shows the variation in the average duration across breaks and their numbers after aggregating contiguous and consecutive movement modes of the same kind into one extended mode.

| **Original** | | | | **Aggregated** | |
| --- | --- | --- | --- | --- | --- |
| **Individual** | **Cluster Width** | **Time** | **Breaks** | **Time** | **Breaks** |
| Agura | 30 | 135.32 | 37 | 223.59 | 22 |
| Akhun | 35 | 115.17 | 6 | 168.50 | 4 |
| Artek | 20 | 115.36 | 102 | 197.57 | 58 |
| Baksan | 35 | 123.91 | 11 | 265.20 | 5 |
| Bardia | 30 | 125.03 | 69 | 182.13 | 46 |
| Borna | 25 | 115.72 | 75 | 159.80 | 51 |
| Borzou | 30 | 121.87 | 71 | 158.91 | 53 |
| Chilmas | 35 | 121.74 | 19 | 160.29 | 14 |
| Elbrus | 30 | 109.30 | 27 | 159.17 | 18 |
| Iran | 30 | 112.27 | 11 | 126.22 | 9 |
| Kaveh | 25 | 111.91 | 76 | 156.31 | 52 |
| Khosta | 30 | 130.46 | 72 | 223.80 | 40 |
| Killi | 30 | 125.71 | 79 | 193.60 | 50 |
| Kodor | 25 | 133.50 | 2 | 133.00 | 2 |
| Laba | 30 | 133.08 | 12 | 194.63 | 8 |
| Laura | 30 | 120.96 | 80 | 222.34 | 41 |
| Leo | 30 | 116.11 | 61 | 160.74 | 42 |
| Tandoureh | 30 | 112.69 | 48 | 163.23 | 31 |
| Victoria | 30 | 127.77 | 64 | 199.60 | 40 |
| Volna | 30 | 127.45 | 64 | 221.39 | 36 |

Table S6. Descriptive statistics on the duration (in hours) of original movement modes for each Persian leopard. Wild leopards were monitored in Iran while rewilded leopards were studied in Russia.

|  |  | **Encamped** | |  |  | **Partial Round-Trip** | | |  | **Ranging** | |  |  | **Round-Trip** | | | | **Wandering** | |  |  |
| --- | --- | --- | --- | --- | --- | --- | --- | --- | --- | --- | --- | --- | --- | --- | --- | --- | --- | --- | --- | --- | --- |
| *Individual* | *Origin* | *n* | *Tot* | *mean* | *sd* | *n* | *Tot* | *mean* | *sd* | *n* | *Tot* | *mean* | *sd* | *n* | *Tot* | *mean* | *sd* | *n* | *Tot* | *mean* | *sd* |
| Agura | Rewilded | 3 | 442.00 | 147.33 | 42.25 | 1 | 82.00 | 82.00 | 0.00 | 21 | 2734.00 | 130.19 | 37.93 | 8 | 1135.00 | 141.88 | 65.54 | 4 | 526.00 | 131.50 | 54.95 |
| Akhun | Rewilded | 2 | 218.00 | 109.00 | 21.21 | 1 | 98.00 | 98.00 | 0.00 | 1 | 134.00 | 134.00 | 0.00 | 2 | 224.00 | 112.00 | 5.66 | 0 | 0.00 | 0.00 | 0.00 |
| Artek | Rewilded | 16 | 1460.00 | 91.25 | 34.26 | 4 | 380.00 | 95.00 | 52.12 | 64 | 8016.00 | 125.25 | 65.36 | 10 | 1034.00 | 103.40 | 30.93 | 7 | 569.00 | 81.29 | 46.49 |
| Baksan | Rewilded | 0 | 0.00 | 0.00 | 0.00 | 1 | 148.00 | 148.00 | 0.00 | 9 | 986.00 | 109.56 | 24.29 | 1 | 192.00 | 192.00 | 0.00 | 0 | 0.00 | 0.00 | 0.00 |
| Bardia | Wild | 37 | 4847.00 | 131.00 | 44.55 | 1 | 170.00 | 170.00 | 0.00 | 7 | 807.00 | 115.29 | 22.32 | 11 | 1246.00 | 113.27 | 33.47 | 12 | 1308.00 | 109.00 | 36.35 |
| Borna | Wild | 25 | 3015.00 | 120.60 | 36.88 | 1 | 148.00 | 148.00 | 0.00 | 24 | 2532.00 | 105.50 | 43.40 | 14 | 1562.00 | 111.57 | 34.78 | 9 | 1105.00 | 122.78 | 44.93 |
| Borzou | Wild | 20 | 2408.00 | 120.40 | 26.27 | 0 | 0.00 | 0.00 | 0.00 | 29 | 3405.00 | 117.41 | 30.67 | 10 | 1255.00 | 125.50 | 29.61 | 12 | 1448.00 | 120.67 | 47.14 |
| Chilmas | Rewilded | 2 | 216.00 | 108.00 | 42.43 | 2 | 190.00 | 95.00 | 21.21 | 9 | 1260.00 | 140.00 | 39.13 | 2 | 226.00 | 113.00 | 32.53 | 4 | 352.00 | 88.00 | 8.79 |
| Elbrus | Rewilded | 6 | 492.00 | 82.00 | 25.04 | 1 | 96.00 | 96.00 | 0.00 | 15 | 1792.00 | 119.47 | 40.73 | 0 | 0.00 | 0.00 | 0.00 | 5 | 485.00 | 97.00 | 36.92 |
| Iran | Wild | 3 | 332.00 | 110.67 | 36.35 | 0 | 0.00 | 0.00 | 0.00 | 4 | 450.00 | 112.50 | 35.98 | 1 | 118.00 | 118.00 | 0.00 | 2 | 236.00 | 118.00 | 5.66 |
| Kaveh | Wild | 17 | 2133.00 | 125.47 | 46.03 | 1 | 100.00 | 100.00 | 0.00 | 32 | 3307.00 | 103.34 | 29.08 | 13 | 1409.00 | 108.38 | 34.69 | 11 | 1179.00 | 107.18 | 48.98 |
| Khosta | Rewilded | 8 | 1064.00 | 133.00 | 35.78 | 2 | 409.00 | 204.50 | 85.56 | 44 | 5346.00 | 121.50 | 40.34 | 8 | 1144.00 | 143.00 | 51.77 | 9 | 1113.00 | 123.67 | 55.49 |
| Killi | Rewilded | 11 | 1459.00 | 132.64 | 57.11 | 2 | 196.00 | 98.00 | 0.00 | 47 | 5639.00 | 119.98 | 48.98 | 13 | 1468.00 | 112.92 | 36.75 | 6 | 918.00 | 153.00 | 20.58 |
| Kodor | Rewilded | 1 | 94.00 | 94.00 | 0.00 | 0 | 0.00 | 0.00 | 0.00 | 0 | 0.00 | 0.00 | 0.00 | 1 | 172.00 | 172.00 | 0.00 | 0 | 0.00 | 0.00 | 0.00 |
| Laba | Rewilded | 5 | 645.00 | 129.00 | 32.66 | 0 | 0.00 | 0.00 | 0.00 | 1 | 102.00 | 102.00 | 0.00 | 3 | 414.00 | 138.00 | 41.90 | 3 | 396.00 | 132.00 | 28.35 |
| Laura | Rewilded | 6 | 714.00 | 119.00 | 50.88 | 4 | 410.00 | 102.50 | 12.58 | 51 | 5966.00 | 116.98 | 37.65 | 9 | 1188.00 | 132.00 | 49.59 | 10 | 1138.00 | 113.80 | 48.96 |
| Leo | Rewilded | 14 | 1392.00 | 99.43 | 34.68 | 2 | 240.00 | 120.00 | 31.11 | 34 | 3913.00 | 115.09 | 38.49 | 4 | 522.00 | 130.50 | 27.15 | 5 | 684.00 | 136.80 | 68.55 |
| Tandoureh | Wild | 23 | 2793.00 | 121.43 | 33.11 | 0 | 0.00 | 0.00 | 0.00 | 10 | 972.00 | 97.20 | 29.13 | 10 | 1037.00 | 103.70 | 38.68 | 5 | 516.00 | 103.20 | 5.93 |
| Victoria | Rewilded | 12 | 1760.00 | 146.67 | 51.18 | 2 | 292.00 | 146.00 | 8.49 | 30 | 3586.00 | 119.53 | 37.79 | 13 | 1494.00 | 114.92 | 54.08 | 7 | 852.00 | 121.71 | 41.19 |
| Volna | Rewilded | 19 | 2295.00 | 120.79 | 40.54 | 0 | 0.00 | 0.00 | 0.00 | 36 | 4235.00 | 117.64 | 60.25 | 8 | 1370.00 | 171.25 | 74.99 | 1 | 70.00 | 70.00 | 0.00 |
| Tot/average |  | 230 | 1388.95 | 112.08 | 34.56 | 25 | 147.95 | 85.15 | 10.55 | 468 | 2759.10 | 111.12 | 33.08 | 141 | 860.50 | 122.86 | 32.11 | 112 | 644.75 | 96.48 | 29.96 |

Table S7. Summary statistics of the duration (in hours), total distance walked (in meters) net squared displacement (NSD, in Km) and number of breaks (N) for each movement mode in the two types of leopards. Wild leopards were monitored in Iran while rewilded leopards were studied in Russia. Ratios refer to the proportion of a metric for a given movement mode out of the total for a given country (only successfully classified modes considered). These figures are derived from the original movement modes.

|  |  |  | **Duration** | | | | **Distance** | | | | **NSDkm** | | | |
| --- | --- | --- | --- | --- | --- | --- | --- | --- | --- | --- | --- | --- | --- | --- |
| *Mode* | *Origin* | *N* | *sum* | *mean* | *sd* | *Ratio* | *sum* | *mean* | *sd* | *Ratio* | *sum* | *mean* | *sd* | *Ratio* |
| encamped | Wild | 125 | 15528.00 | 124.22 | 38.20 | 0.39 | 5421765.42 | 43374.12 | 20500.57 | 0.40 | 1778.92 | 14.23 | 19.10 | 0.15 |
| encamped | Rewilded | 105 | 12251.00 | 116.68 | 44.02 | 0.16 | 2313089.67 | 22029.43 | 18254.72 | 0.15 | 1143.47 | 10.89 | 25.55 | 0.02 |
| partial round-trip | Wild | 3 | 418.00 | 139.33 | 35.80 | 0.01 | 114719.29 | 38239.76 | 14175.77 | 0.01 | 48.14 | 16.05 | 11.31 | 0.00 |
| partial round-trip | Rewilded | 22 | 2541.00 | 115.50 | 44.64 | 0.03 | 533319.48 | 24241.79 | 15406.45 | 0.03 | 1572.74 | 71.49 | 116.32 | 0.02 |
| ranging | Wild | 106 | 11473.00 | 108.24 | 33.15 | 0.29 | 3952453.66 | 37287.30 | 22775.79 | 0.29 | 7751.95 | 73.13 | 76.96 | 0.66 |
| ranging | Rewilded | 362 | 43709.00 | 120.74 | 47.58 | 0.57 | 8885053.44 | 24544.35 | 15885.83 | 0.57 | 57440.91 | 158.68 | 235.16 | 0.81 |
| round-trip | Wild | 59 | 6627.00 | 112.32 | 33.59 | 0.17 | 2172852.34 | 36828.01 | 19023.41 | 0.16 | 731.36 | 12.40 | 18.15 | 0.06 |
| round-trip | Rewilded | 82 | 10583.00 | 129.06 | 51.10 | 0.14 | 2345838.98 | 28607.79 | 17835.38 | 0.15 | 2406.22 | 29.34 | 68.49 | 0.03 |
| wandering | Wild | 51 | 5792.00 | 113.57 | 40.48 | 0.15 | 1915136.43 | 37551.69 | 19053.53 | 0.14 | 1519.59 | 29.80 | 46.56 | 0.13 |
| wandering | Rewilded | 61 | 7103.00 | 116.44 | 47.44 | 0.09 | 1475092.76 | 24181.85 | 19298.37 | 0.09 | 8779.95 | 143.93 | 374.62 | 0.12 |

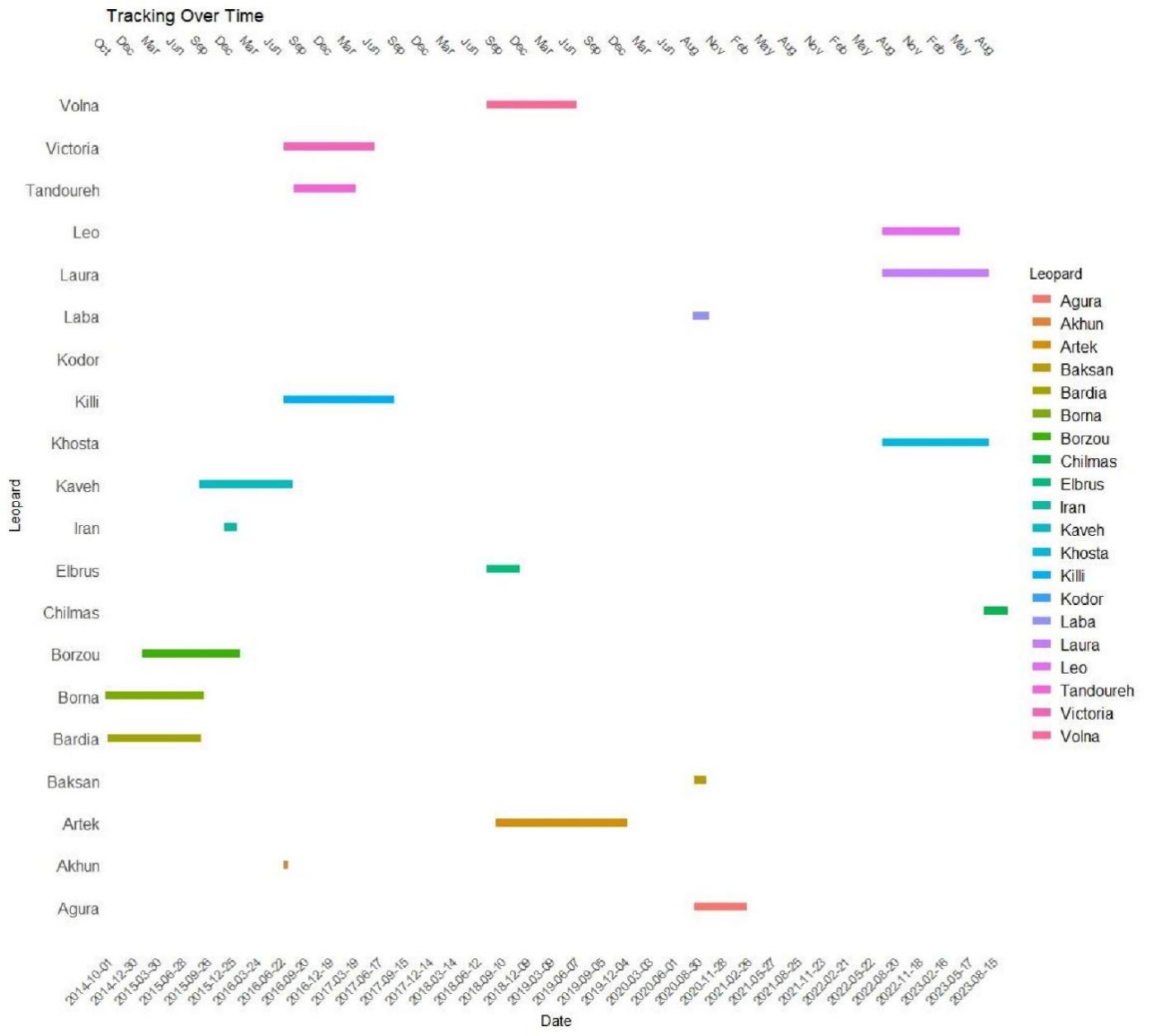

Figure S1. Tracking duration for all Persian leopards included in the study.

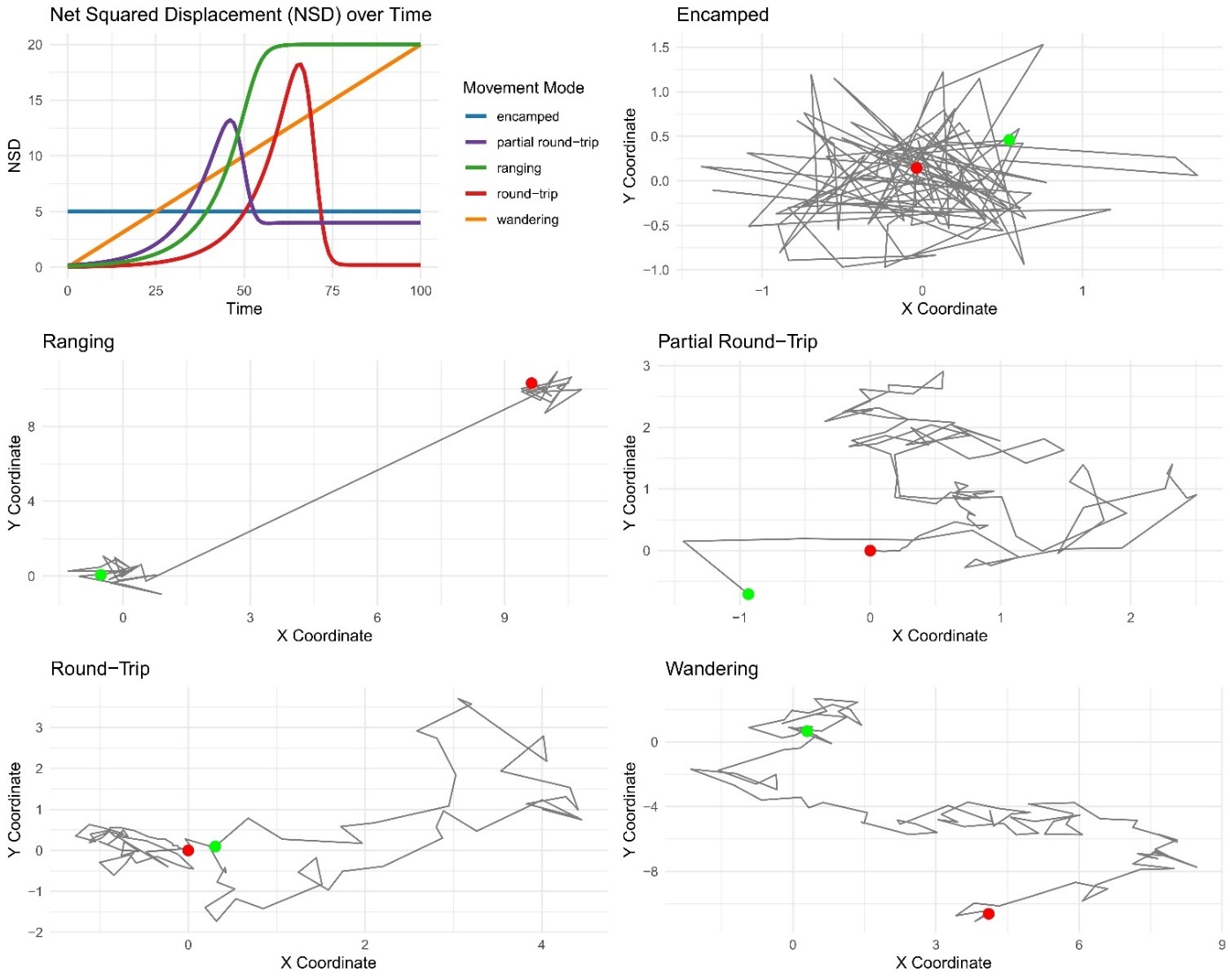

Figure S2. Linear and non-linear curves applied to Net Squared Displacement (NSD) patterns post-BCPA segmentation analysis. For the mathematical details of these curves, refer to Morelle et al. (2017). The figure also categorizes movement patterns into respective modes alongside the curves.

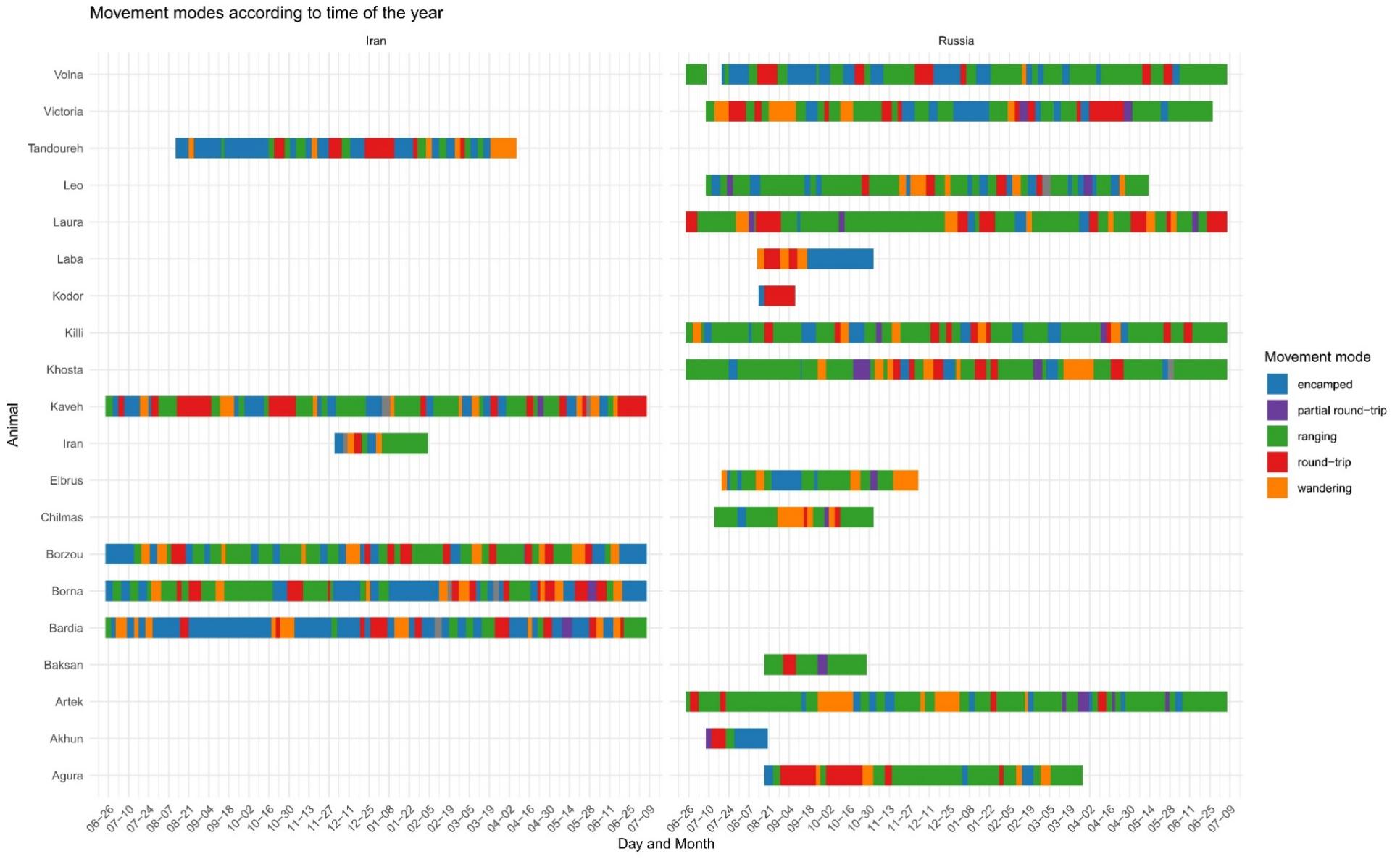

Figure S3. Movement modes of each Persian leopard by the time of year and country. Unclassified segments, not featured in the legend, are marked in grey.

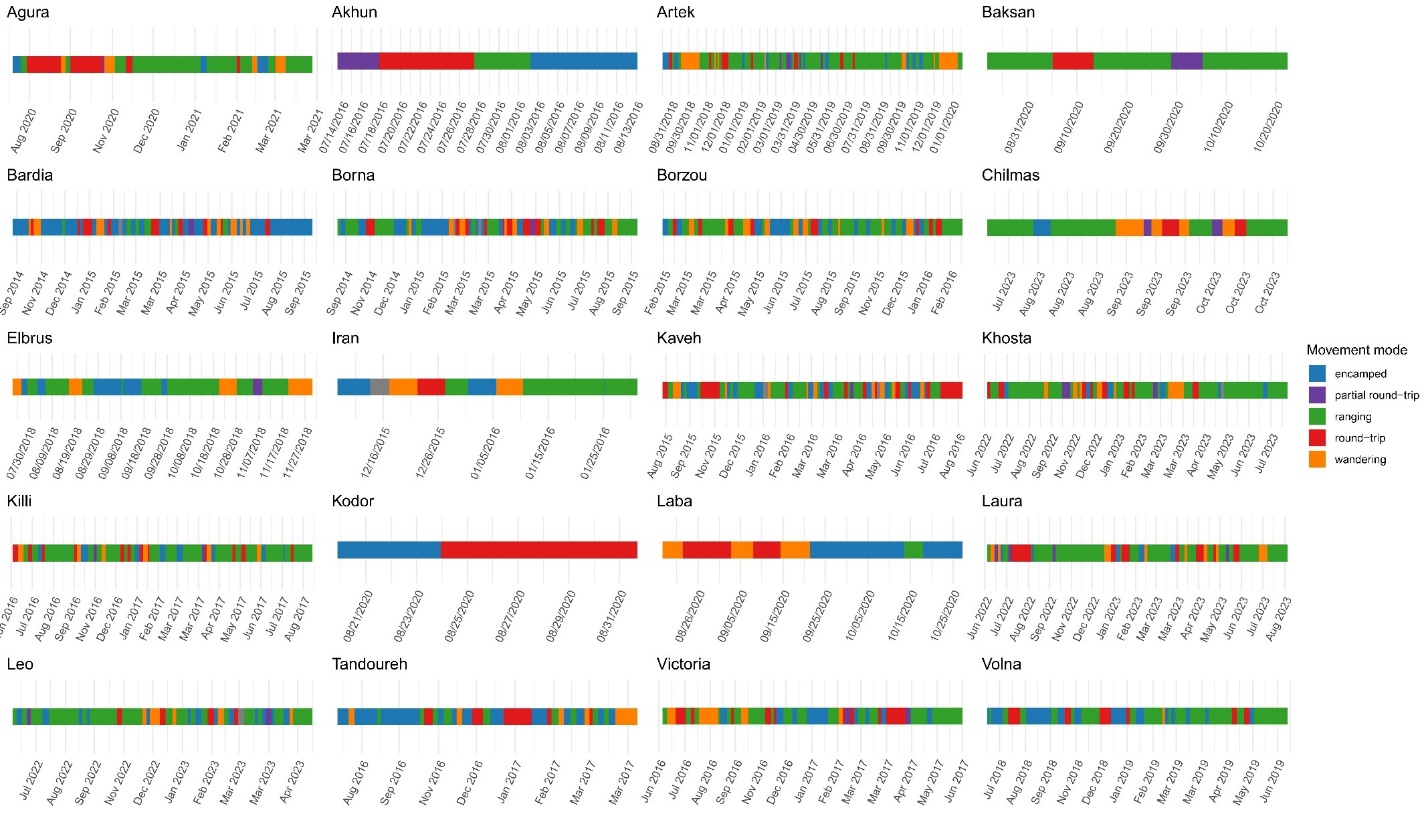

Figure S4. Movement modes identified in each Persian leopard's track. Unclassified segments, not depicted in the legend, are represented in grey.

### References

Bjørneraas, K., B. Van Moorter, C. M. Rolandsen, and I. Herfindal. 2010. Screening global positioning system location data for errors using animal movement characteristics. The Journal of Wildlife Management 74:1361–1366.

Calabrese, J. M., C. H. Fleming, and E. Gurarie. 2016. ctmm: an R package for analyzing animal relocation data as a continuous‐time stochastic process. Methods in Ecology and Evolution 7:1124–1132.

Farhadinia, M. S., P. J. Johnson, D. W. Macdonald, and L. T. B. Hunter. 2018. Anchoring and adjusting amidst humans: Ranging behavior of Persian leopards along the Iran-Turkmenistan borderland. PLoS ONE 13.

Farhadinia, M. S., M. Kaboli, M. Karami, and H. Farahmand. 2014. Patterns of sexual dimorphism in the Persian Leopard (Panthera pardus saxicolor) and implications for sex differentiation. Zoology in the Middle East 60:195–207.

Farhadinia, M. S., I. Memarian, K. Hobeali, A. Shahrdari, B. Ekrami, J. Kaandorp, and D. W. Macdonald. 2017. GPS collars reveal transboundary movements by Persian leopards in Iran. Cat News 65:28–30.

Gupte, P. R., C. E. Beardsworth, O. Spiegel, E. Lourie, S. Toledo, R. Nathan, and A. I. Bijleveld. 2022. A guide to pre‐processing high‐throughput animal tracking data. Journal of Animal Ecology 91:287–307.

Morelle, K., N. Bunnefeld, P. Lejeune, and S. A. Oswald. 2017. From animal tracks to fine‐scale movement modes: a straightforward approach for identifying multiple spatial movement patterns. Methods in Ecology and Evolution 8:1488–1498.

Rozhnov, V., A. Yachmennikova, N. Dronova, A. Pkhitikov, M. Magomedov, I. Chestin, and Alibekov. AB. 2020. The restoration  of Persian leopard in the Caucasus (scientific  approach). KMK Scientific Press Ltd., Moscow.
